# Correlation guided Network Integration (CoNI) reveals novel genetic regulators of hepatic metabolism

**DOI:** 10.1101/2020.01.29.924944

**Authors:** Valentina S. Klaus, Sonja C. Schriever, Andreas Peter, José Manuel Monroy Kuhn, Martin Irmler, Janina Tokarz, Cornelia Prehn, Gabi Kastenmüller, Johannes Beckers, Jerzy Adamski, Alfred Königsrainer, Timo D. Müller, Martin Heni, Matthias H. Tschöp, Paul T. Pfluger, Dominik Lutter

## Abstract

The steadily increasing amount of newly generated omics data of various types from genomics to metabolomics is a chance and a challenge to systems biology. To fully use its potential, one key is the meaningful integration of different types of omics. We here present a fully unsupervised and versatile correlation-based method, termed Correlation guided Network Integration (CoNI), to integrate multi-omics data into a hypergraph structure that allows for identification of effective regulators. Our approach further unravels single transcripts mapped to specific densely connected metabolic sub-graphs or pathways. By applying our method on transcriptomics and metabolomics data from murine livers under standard chow or high-fat-diet, we isolated eleven genes with a regulatory effect on hepatic metabolism. Subsequent *in vitro* and *ex vivo* experiments in human liver cells and human obtained liver biopsies validated seven candidates including *INHBE* and *COBLL1*, to alter lipid metabolism and to correlate with diabetes related traits such as overweight, hepatic fat content and insulin resistance (HOMA-IR). Last, we successfully applied our methods to an independent data-set to confirm its versatile and transferable character.

## INTRODUCTION

In the era of systems biology and multi-omics high throughput data generation, there is an unmet need for effective, versatile and straightforward tools and approaches to compare and integrate large and complex data sets built on a plurality of different genetic classifiers and types of biomolecules with complex inter- and intra-regulatory connections [1, 2]. In particular, for capturing genetic mechanisms associated with metabolic disorders, which typically affect multiple layers of biological regulations, effective tools are needed to further deepen our understanding on the control of metabolism. Current integration approaches to interrogate complex metabolic networks [3] are largely built on integrating genetic information via associative approaches or via utilizing additional prior knowledge. These attempts to integrate multi-omics data combine metabolic profiles with genetic information, i.e. genetic variants or direct correlations of transcripts with metabolites [4, 5], use prior knowledge to map genes and enzymes on metabolic pathways [6-8] or create deterministic models that abstract enzymatic reactions in metabolic pathways using gene or enzyme levels as rate-limiting denominators [9-11].

However, at current such attempts to integrate genetic information into metabolic networks lack a suitable underlying model that reflects the relations between single metabolites driven via regulatory genes directly or indirectly affecting enzymatic reactions in metabolic pathways. A further limiting factor is that metabolome methods typically only capture a few hundred to thousand different metabolites of estimated more than 40,000 metabolites [12], which dramatically limits the use of prior knowledge. Accordingly, novel approaches to reveal distinct regulatory genes are warranted.

One area where sophisticated multi-omics data integration could be particularly useful is the study of hepatic steatosis, a pathological state of the liver that is characterized by excess fat accumulation due to impaired lipid metabolism [13]. Understanding the molecular mechanisms that drive the accumulation of lipids in the liver under perturbed metabolic homeostasis could be key to new targets for early diagnosis and treatment of hepatic steatosis.

In this study, we present a novel statistical method for correlation-based network integration (CoNI) of generic character and conceivable for multiple approaches. We applied CoNI to murine liver metabolome and transcriptome data to unravel previously hidden gene-metabolite interactions that exert major changes to hepatic metabolite levels under normal dietary conditions and under obesogenic stress. These CoNI analyses generated two independent networks for chow-fed lean vs. high fat diet (HFD) fed obese mice, each mirroring the respective genetic-metabolic interactome. Both networks were then evaluated and compared based on graph-theoretic properties, differential expression patterns and the presence of single-nucleotide polymorphisms (SNPs) associated with obesity and type 2 diabetes in humans. Based on these criteria, eleven genes were considered as most promising candidates with likely metabolic relevance, and subjected to validation by siRNA knockdown *in vitro* and transcriptional profiling of human liver biopsies *ex vivo*. Of those eleven candidate genes, seven had significant effects on cellular metabolite levels and/or were correlated with hepatic triglyceride levels, BMI or insulin resistance in human liver specimens. Among these, four had already been linked to hepatic lipid metabolism, whereas three have not been described in the context of high-fat diet feeding, hepatosteatosis, obesity and/or diabetes. Overall, these data demonstrate that our novel CoNI framework is a fully data-driven, flexible and versatile tool for multiple omics data integration and gives way to a meaningful interpretation.

## RESULTS

### 1. CoNI: Correlation guided Network Integration

The CoNI framework utilizes correlations and partial correlations to combine two types of omics data **(driver data, linked data)**, thereby generating a hypergraph where edges are formed by the driver data specifying the impact onto the interaction of the linked data. Here, CoNI was applied to integrate hepatic transcriptome (driver) data with hepatic metabolome (linked) data from chow-fed lean and HFD-fed obese mice. CoNI is built on the general concept (Figure 1) that Pearson correlations are calculated for each pair of metabolites followed by removing the linear effect of each gene on this correlation by calculating the partial correlation coefficient. For *K* genes and *M* metabolites, we thus generate *K M*x*M* matrices containing the partial correlation coefficients. Next, an adapted Steiger test [14] is applied to compute the impact of each gene on the metabolite correlation, thereby generating *K* adjacency matrices, one for each gene. From these adjacency matrices, edges significantly connecting nodes with a p-value below a predefined alpha (p < 0.05) are selected and assembled in form of an integrated graph where the nodes refer to the metabolite pairs and the edges to the controlling genes. A gene can thereby be mapped to multiple edges, and edges may consist of multiple genes. Finally, this network assembly is used to identify local regulator genes (LRGs), i.e. genes locally enriched in a densely connected sub-graph. In our data, these LRGs unravel previously hidden genes that regulate specific metabolic sub-graphs and thus gene-metabolite sub-networks. We also successfully applied CoNI to integrate proteomics into a lipidomics network of lungs of C57BL/6 mice under fresh air and smoking conditions (see SI).

**Figure 1:**
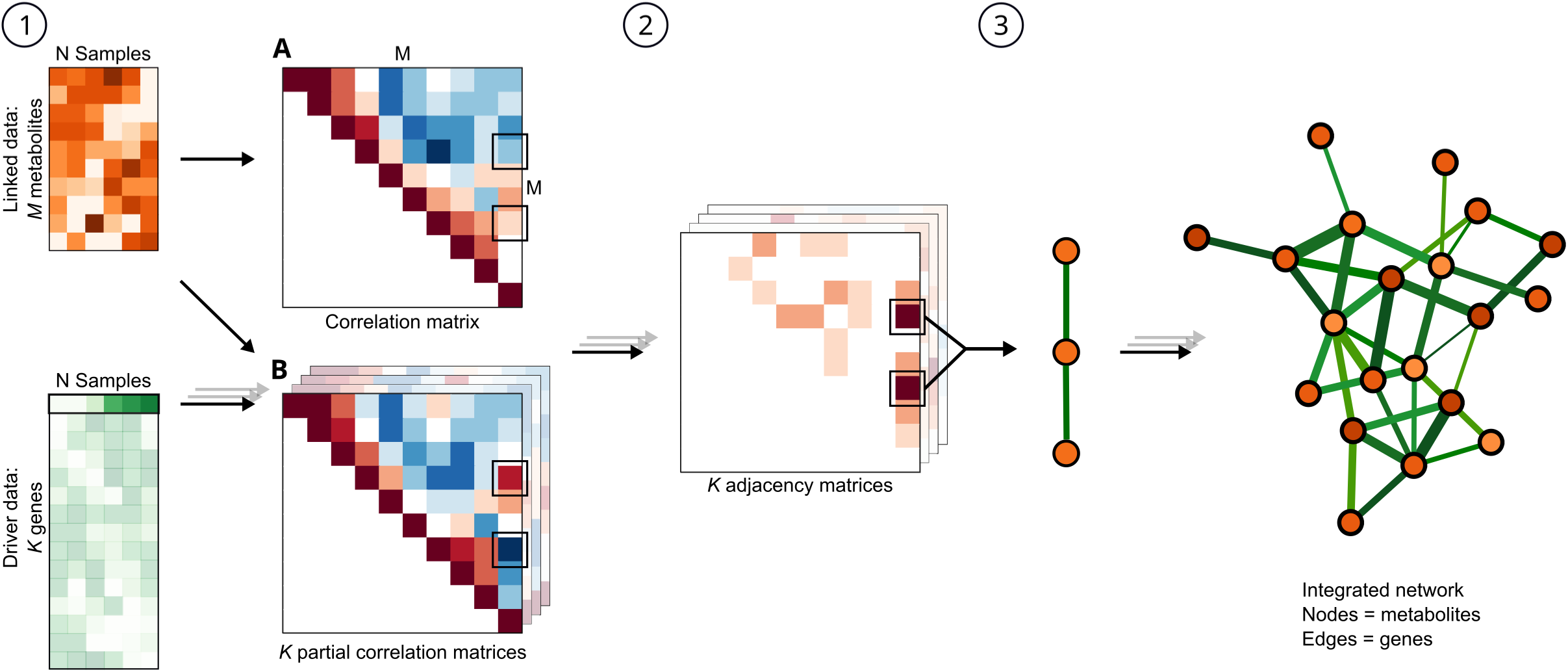
CoNI workflow. **1)** Calculation of a full pairwise correlation matrix (**A**) and partial correlation analysis combining the metabolite concentrations with the transcript expression profiles; for each pair of metabolites partial correlation scores are computed by subtracting an estimated regulatory effect for each gene *k*. This results in *K* partial correlation matrices (**B**). **2)** Calculation of *K* adjacency matrices by selecting metabolite pairs significantly altered by individual genes. **3)** Selection of significant triplets (metabolite pair plus gene) and construction of an undirected, weighted graph with correlated metabolite pairs as nodes and influencing genes setting up the edges. The number of genes connecting the two respective metabolite nodes determines the edge weight, indicated by thickening of the line.

### 2. Transcriptional and metabolic profiling of livers from chow and HFD-fed mice

In order to investigate effects of diet-induced obesity (DIO) on the liver transcriptome and metabolome, 10 male C57Bl6J mice were exposed either to standard chow or to 58% HFD (D13331, Research Diets) for 22 weeks, respectively. Exposure to the high-caloric diet resulted in a significantly higher body weight (BW, Chow 33.5 g ± 1.6 g; HFD 49.2 g ± 4.5 g, p < 0.0001, mean ± SD) (Figure 2A). Two mice from the HFD cohort with body weights comparably lower to those of the other HFD-fed mice (BW at week 22: 37.5 g and 33.3 g) were excluded from all further analyses. HFD-fed obese mice showed increased plasma triglyceride and cholesterol levels (Figure 2B,C), were hyperinsulinemic (Figure 2D) and had increased hepatic triglyceride (TAG) stores (Figure 2E, Table 1) compared to chow-fed lean controls.

**Table 1.** Characteristics of the mouse cohorts (mean ± SD).

|  | Chow (n=10) | HFD (n=8) | p-value |
| --- | --- | --- | --- |
| Body weight (end) (g) | 33.49 ± 1.56 | 49.2 ± 4.49 | < 0.0001 |
| Liver triglyceride levels (μg/mg tissue) | 4.13 ± 0.9 | 9.4 ± 3.11 | 0.0001 |
| Plasma triglyceride levels (mg/dl) | 44.09 ± 6.84 | 54.49 ± 8.43 | 0.0105 |
| Plasma cholesterol levels (mg/dl) | 45.74 ± 6.84 | 121.24 ± 18.78 | <0.0001 |
| Plasma insulin levels (ng/ml) | 1.58 ± 0.85 | 7.32 ± 5.65 | 0.0057 |

**Figure 2:**
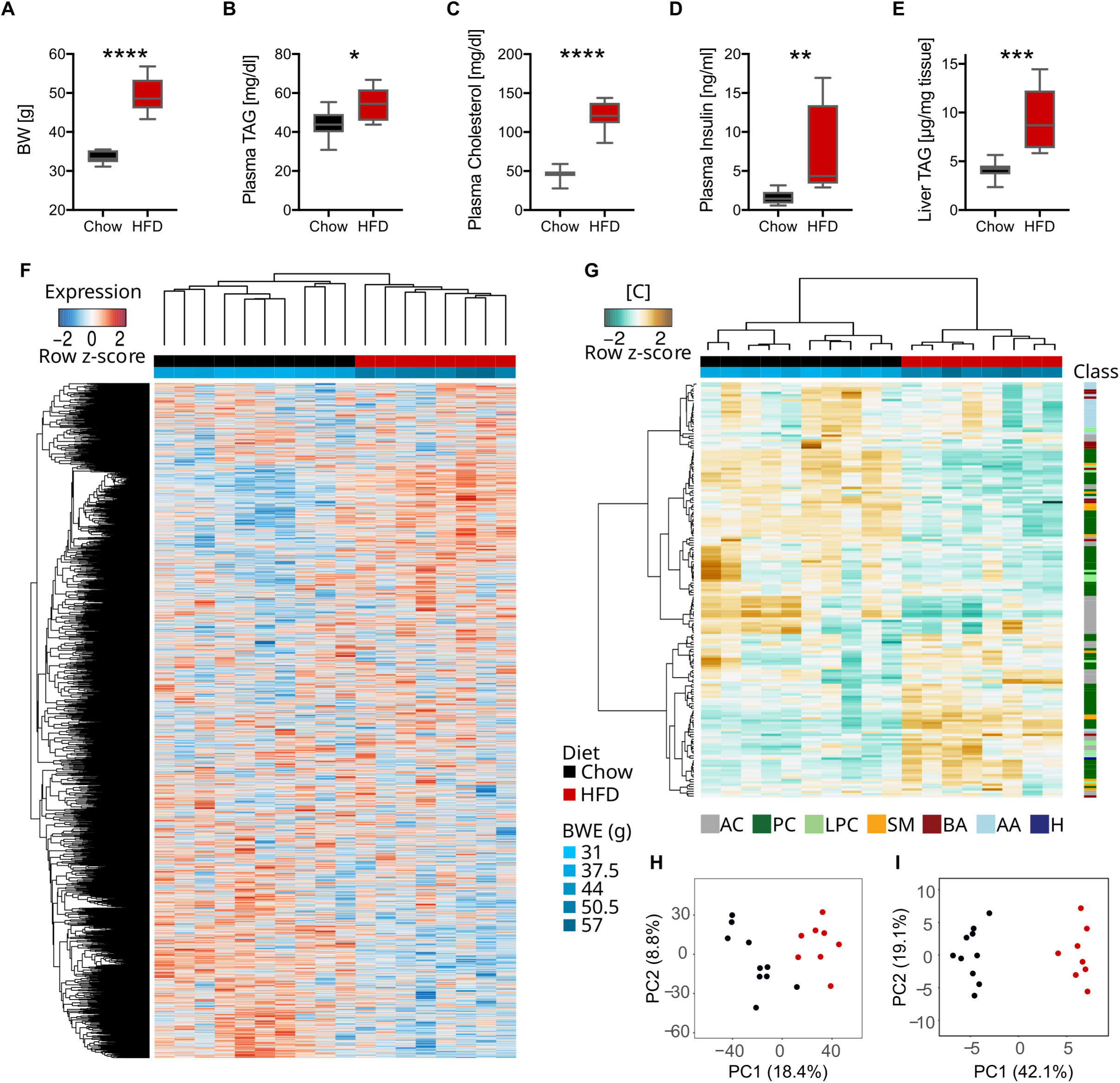
Transcriptional and metabolic profiling of murine livers under normal conditions and obesogenic stress. Barplots comparing mice after 22 weeks of chow or HFD for **(A)** body weight, **(B)** plasma triacylglyceride (TAG) and plasma cholesterol **(C)**, plasma insulin **(D)**, and hepatic TAG levels **(E)**. Asterisks indicate the significance of the differences between the factors (*p <= 0.05; **p <= 0.01; ***p <= 0.001). Errorbars show standard error of the mean (SEM) **(F)** Heatmap with 10,159 hepatic mRNAs transcripts detected in chow-fed (black color, upper bar) and HFD-fed (red color, upper bar) mice. The lower color bar indicates individual body weights measured at the end of the study (BWE). **(G)** Heatmap with concentrations of 175 detected metabolites. Metabolite classes are indicated in the right color bar: amino acids (AA), sphingomyelins (SM), lyso-phosphatidylcholines (LPC), phosphatidylcholines (PC), biogenic amines (BA), acylcarnitines (AC), hexose (H). PCA plot of transcript expression **(H)** and metabolite concentrations **(I)** for chow (black) and HFD (red). The amounts of variance explaining the differences are given in brackets.

Next, hepatic metabolism was analyzed by transcriptional and metabolic profiling using Affymetrix Microarrays and the Absolute*IDQ*™ p180 Kit resulting in a total of 10,159 genes (Figure 2F) expressed in all mouse livers and a total of 175 detected metabolites (Figure 2G, Table S1) split into seven classes: 40 acylcarnitines (AC), 76 phosphatidylcholines (PC), 14 lyso-phosphatidylcholines (LPC), 12 sphingomyelins (SM), 12 biogenic amines (BA), 20 amino acids (AA) and 1 hexose (H). In total, we identified 989 genes as significantly differentially expressed between chow and HFD from which 501 were up-regulated on HFD and 488 down-regulated, respectively (Figure S1A, Table S2). Functional enrichment analyses based on Gene Ontology (GO) of the up- and down-regulated genes revealed numerous metabolic and in particular lipid-related processes (Figure S1B, Table S2). The most significantly regulated genes were *Cyp2b9*, which is involved in hepatic lipid homeostasis [15], and *Ces2a*, which controls a hepatic lipid network dysregulated in obese humans and mice [16].

We also identified 91 metabolites as significantly altered by diet from which 55 were up-regulated on HFD and 36 were down-regulated (Figure S1C, Table S1). The most prominently regulated metabolite classes were the SM (67% regulated), followed by PC (64%) and the AC (45%). Principal component analyses (PCA) identified diet as the main contributor to explain variance in gene expression (Figure 2H) as well as the main driver of metabolite variance (Figure 2I).

### 3. Correlation maps of diet-altered metabolites

To further investigate diet-induced changes on liver metabolism, we generated two diet-dependent correlation maps by calculating the pairwise Pearson correlation coefficients of all metabolites (Figure 3A). For both diets, we observe a slight negative-skew of the correlation coefficient distribution, which was also observed by others [17] (Figure S2A). We identified 2,488 significantly correlated metabolite pairs for chow and 2,322 for HFD with a p < 0.05 (non adjusted), 923 were identified in both diets (Figure 3B). Furthermore, 1,023 (non-adjusted) metabolite pairs were identified that showed a significant change in their correlation from chow to HFD, indicating that the administered diet substantially alters the hepatic metabolism.

**Figure 3:**
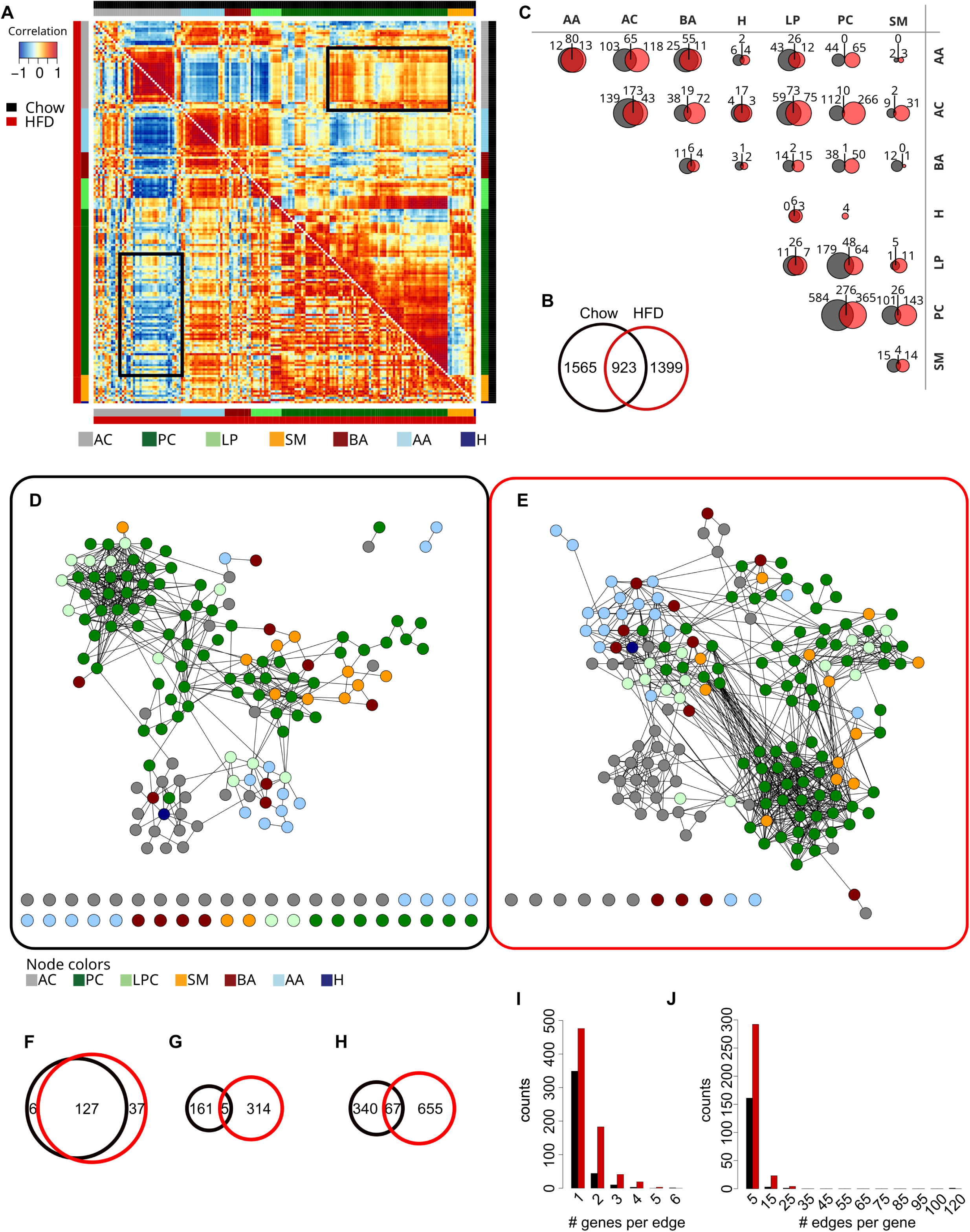
Metabolite correlations under chow and HFD. **(A)** Pairwise metabolite correlation matrix showing the correlation coefficients obtained for chow (black) in the upper right and for HFD (red) in the lower left triangle. The metabolite classes are indicated by the respective color bars. Black boxes mark correlations between AC and PC in chow and HFD. **(B)** Venn diagram showing significantly correlating metabolite pairs differing and overlapping between chow and HFD. **(C)** Metabolite class comparison of significantly correlated metabolite pairs in chow and HFD. The sizes of the Venn diagrams correspond to the number of metabolite pairs. **(D, E)** Integrated graphs generated using CoNI for chow (**D**) and HFD **(E)**. The nodes of the graphs refer to metabolites and edges to genes that significantly affect metabolite correlation. Node colors reflect the metabolite class. Unconnected nodes are displayed at the bottom. **(F)** Comparison of connected metabolite nodes between chow (black) and HFD (red). **(G)** Comparison of genes contained in edges of both graphs. **(H)** Comparison of significantly correlated metabolite pairs between the two graphs. **(I)** Number of genes per edge and **(J)** number of edges per gene in both graphs.

We deepened our investigations and analyzed class composition of the correlated metabolite pairs (Figure 3C). The metabolite class with the maximum change in correlated pairs was SM (Jaccard Index = 0.12). In contrast, AA mainly maintain their correlations (Jaccard Index = 0.76) indicating that amino acids interact independent of dietary conditions. Between classes we generally observed a substantial change in correlations between chow and HFD with Jaccard indices between 0.01 (PC – BA) and 0.71 (H – AC). We further observed a striking difference between AC and PC with the highest absolute numbers of changes driven by the shift from positive correlations under chow to negative ones under HFD (Figure 3A, C, and S2B). For the pairs SM – BA, SM – AA, SM – H, AA – PC and PC – H no overlap of pairs with a significant correlation in both diets could be observed. Taken together, correlation analyses revealed substantial diet-dependent changes in metabolite regulation, which was previously reported in a circadian manner over multiple tissues [18].

### 4. Estimating genetic impact on metabolic networks

We next assessed the genetic impact on the metabolite network formed under each dietary condition by integrating gene expression and metabolite concentration profiles with our CoNI approach. Two independent networks for chow and HFD were generated (Figure 3D, E), constructed of 485 triplets (gene and metabolite pairs) for the chow and 1,058 triplets for the HFD set (Table S3). Of the 175 metabolites used for the analysis 133 were connected in the chow (Figure 3D) and 164 metabolites in the HFD network (Figure 3E), respectively. Of the connected metabolites, we found 127 in both networks (Figure 3F). We next analyzed the numbers of genes forming the edges between the metabolite pairs, and found 166 different genes in the chow network, 319 genes in the HFD network, and only five genes in both networks (Figure 3G). When comparing pairs of connected metabolites, 67 were identical in both diets, 340 were only connected in the chow and 655 only in the HFD network (Figure 3H). Node degree distributions of both networks showed consistent higher degrees for HFD compared to chow diet (Figure S3A), which was also observed when comparing the node degrees for the specific metabolite classes (Figure S3B). In contrast to the elevated but similar distribution of node degrees within the HFD network, the chow network shows a trend towards increased node degrees for PC and LPC compared to the other metabolite classes. A striking characteristic of the inferred networks is that both tend to be organized in communities (Figure S4A, B), or densely connected sub-networks, which mainly refer to metabolite classes but are partly reorganized on dietary change (Figure S4C). This reorganization was also observed in further network characteristics such as the shortest path length (Figure S5).

Analogous to this, the genes connecting metabolite pairs in the two inferred networks massively changed depending on dietary conditions (Figure 3G). Only five genes are present in edges of both graphs (*Gm4553, Hnrnpm, Tap1, Xpo7, Eya3*) from which *Tap1* was the only one differentially expressed between HFD and chow diet (Figure S6A,B). Comparing the number of genes that map to single edges, we observed that most (85.75% in chow, 65.93% in HFD) edges consist of a single gene and the maximum number of genes per edge was six in chow and five in HFD (Figure 3I, Table S3). The distribution of individual genes over the edges (Figure 3J) showed that most genes appeared in five or less edges. The highest distribution of a single gene in both networks was found for *Nop16*, appearing in 113 of 407 edges in the chow and *Cobll1*, appearing in 25 of 722 edges in the HFD network.

To further classify the genes found in both networks a functional enrichment analysis using KEGG pathways and GO biological processes was performed (Table S3). For the chow network, we did not identify informative categories for the individual genes. However, the individual genes of the HFD network were identified to be enriched in the KEGG categories ‘glycerolipid metabolism’ and ‘non-alcoholic fatty liver disease’ (NAFLD). This finding confirms that our CoNI approach was reflecting the metabolic phenotype of the HFD group as the mice showed elevated TAG levels in the liver (Figure 2E), a hallmark of fatty liver disease.

### 5. Effective network genes

Our primary motivation of integrating transcriptional data into a metabolic network was to identify differences in the genetic programs that control hepatic metabolism under healthy conditions vs. acute hepatic steatosis. Thus, we defined local regulator genes (LRGs) as genes significantly enriched within a local sub-graph (see methods), thereby controlling a densely connected metabolic sub-network. This resulted in 20 identified LRGs in the chow and 59 in the HFD network, with no overlap (Table S3). From these 79 LRGs eight were differentially expressed, one from the chow network (*Ddx3x*) and seven from the HFD network, namely *Myc, Arhgap24, Smim13, Rapgef4, Cd82, Inhbe*, and *Gk* (Figure 4A-H, S7, Table 2, S3). The isolated sub-networks of the selected LRGs comprise between six (*Ddx3x, Smim13*) and 14 metabolites (*Cobll1*).

**Table 2.**
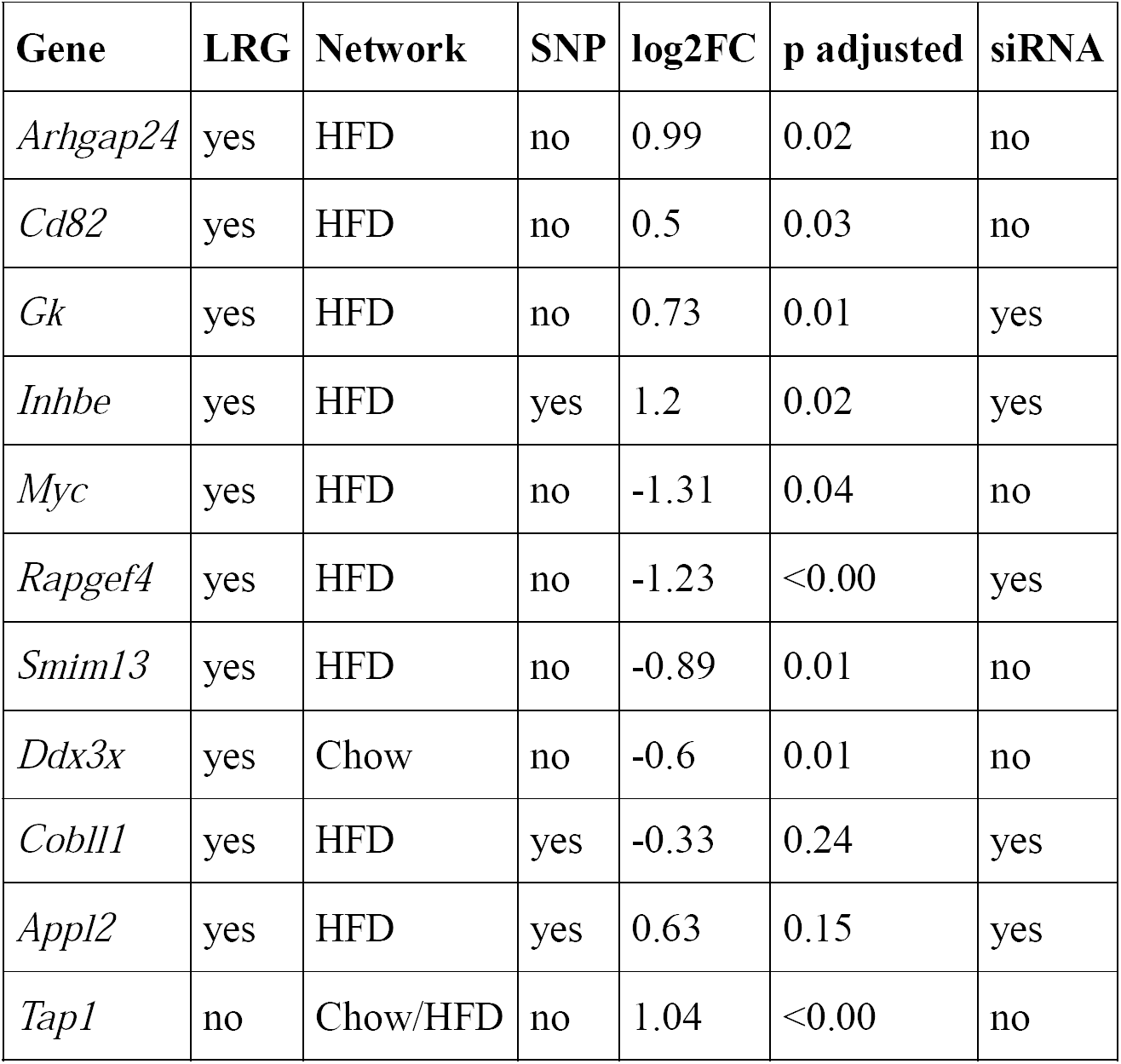
Characteristics and selection criteria for the eleven genes further subjected to validation experiments: identification as LRG, present in specific network, identification as obesity and type 2 diabetes related genetic variant (SNP), differential hepatic expression between chow and HFD mice (log2FC and p) and selection for siRNA KD experiments.

| Gene | LRG | Network | SNP | log2FC | p adjusted | siRNA |
| --- | --- | --- | --- | --- | --- | --- |
| *Arhgap24* | yes | HFD | no | 0.99 | 0.02 | no |
| *Cd82* | yes | HFD | no | 0.5 | 0.03 | no |
| *Gk* | yes | HFD | no | 0.73 | 0.01 | yes |
| *Inhbe* | yes | HFD | yes | 1.2 | 0.02 | yes |
| *Myc* | yes | HFD | no | -1.31 | 0.04 | no |
| *Rapgef4* | yes | HFD | no | -1.23 | <0.00 | yes |
| *Smim13* | yes | HFD | no | -0.89 | 0.01 | no |
| *Ddx3x* | yes | Chow | no | -0.6 | 0.01 | no |
| *Cobll1* | yes | HFD | yes | -0.33 | 0.24 | yes |
| *Appl2* | yes | HFD | yes | 0.63 | 0.15 | yes |
| *Tap1* | no | Chow/HFD | no | 1.04 | <0.00 | no |

**Figure 4:**
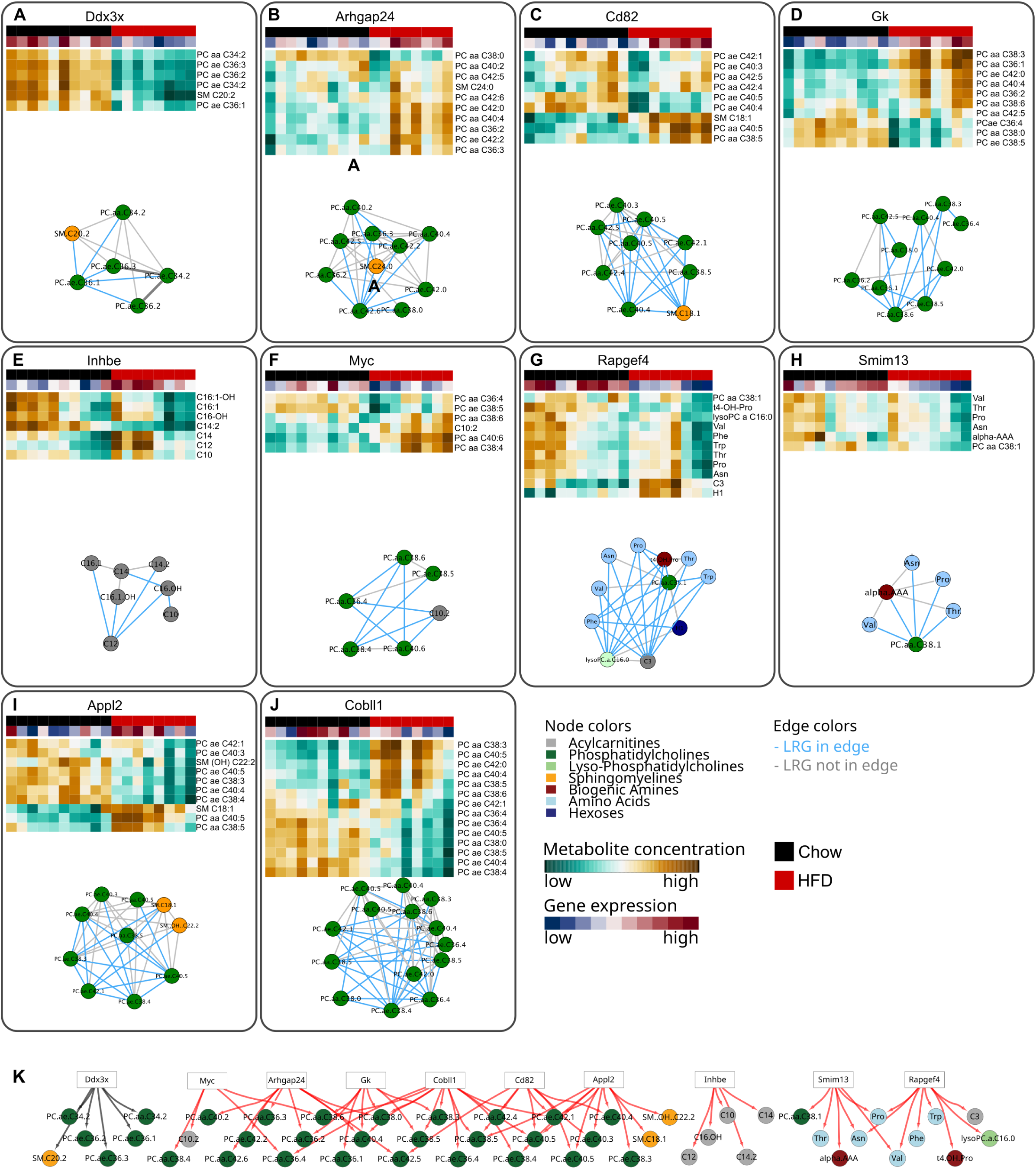
Local regulator genes (LRGs) and associated metabolite sub-graphs. **(A-J)** metabolite sub-graphs for the selected LRGs. Node colors refer to the different metabolite classes; the respective heatmaps show the concentrations of node metabolites. Gene expression levels of the respective LRG displayed in the upper color bar for chow (black) and HFD (red). Blue edges denote edges that contain the LRG in the sub-graph, grey edges are void of the LRG. LRGs were identified in the chow **(A)** and HFD **(B-J)** network from which eight were differentially expressed dependent on diet (**A-H**) and for three genes SNPs associated with obesity and related disease markers could be identified (**E**,**I**,**J**). **(K)** Combined LRG network. Edges display direct gene – metabolite connections. Edge colors refer to diet, i.e. chow (black) or HFD (red).

Additionally, we queried the Type 2 Diabetes Knowledge Portal [19] to determine LRGs that contain Single Nucleotide Polymorphisms (SNPs) associated with obesity and related disease markers. For our set of 79 LRGs, we could identify SNPs for three LRGs of the chow network and for 17 LRGs of the HFD network (Table S4, SI). Among the genes most prominently associated with SNPs were *Appl2* (Figure 4I), which mediates insulin signaling, endosomal trafficking, adiponectin and other signaling pathways [20] and *Cobll1* (Figure 4J), which was strongly associated with several obesity and type 2 diabetes related markers [21, 22]. We further found the differentially expressed LRG *Inhbe*, a hepatokine which was recently linked to insulin resistance in human livers [23]. For all selected LRGs, the regulated metabolites show substantial differences between chow and HFD (Figure 4A-J).

To test for interrelation of the selected LRGs we next combined the isolated metabolite-gene sub-networks and obtained three interconnected sub-graphs for HFD and one for chow (Figure 4K). The largest interconnected sub-graph of the HFD network contained mostly PC and was controlled by six genes. In contrast, the HFD network derived LRGs *Smim13* and *Rapgef4* included BA, PC, LPC and AC metabolites, but were interconnected mainly by an overlapping mixture of AA. The third HFD sub-network was under the control of the LRG *Inhbe* and contained only AC.

### 6. Network genes are regulators of human hepatic metabolism

To assess the translational relevance of these findings derived from our novel CoNI methodology, we selected eleven candidate genes for validation in human liver biopsies. Ten genes were locally enriched in one of the networks (LRGs), eight of these showed differential expression between diets, three were associated with human obesity-related SNPs, and one non-LRG (*Tap1*) was differentially expressed and present in both networks (Table 2 and SI).

The eleven candidate genes were quantified by qPCR analyses for hepatic mRNA expression levels in liver biopsies of 170 patients. Expression levels were then correlated with liver fat content and BMI (Figure S8). Anthropometrics and metabolic characteristics of these subjects, which covered a wide range of hepatic triglyceride content, are shown in Tables S5A, B. Associations of hepatic mRNA levels with insulin resistance (HOMA-IR) were additionally analyzed for a subgroup of 77 subjects where fasting blood samples were available. Significant associations of gene expression and metabolic traits were found for five of the eleven genes (Figure 5A-K, Table 3). *GK, INHBE, TAP1* and *MYC* were associated with BMI (Figure 5A,B,C,E), whereas *INHBE* and *SMIM13* expression correlated significantly with liver fat content (Figure 5G,I). Strikingly, the expression of the LRG *INHBE* in human liver was not only significantly associated with BMI and liver TAG content but also with the HOMA-IR (Figure 5B,G,K).

**Table 3.** Associations between hepatic biopsy gene expression levels and metabolic traits BMI (N=170), hepatic TAG (N=170) and HOMA-IR (N = 77) in human liver samples.

| Name | BMI p | BMI r<sup>2</sup> | TAG p | TAG r<sup>2</sup> | HOMA-IR<br>p | HOMA-IR r<sup>2</sup> |
| --- | --- | --- | --- | --- | --- | --- |
| ***RAPGEF4*** | 0.7570 | 0.0006 | 0.7010 | 0.0009 | 0.2330 | 0.0189 |
| ***ARHGAP24*** | 0.0934 | 0.0167 | 0.0899 | 0.0171 | 0.8610 | 0.0004 |
| ***GK*** | **0.0034** | 0.0500 | 0.0899 | 0.0171 | 0.1010 | 0.0355 |
| ***COBLL1*** | 0.7430 | 0.0006 | 0.1570 | 0.0119 | 0.7300 | 0.0016 |
| ***INHBE*** | **0.0001** | 0.0842 | **< 0.0001** | 0.0990 | **0.0088** | 0.0906 |
| ***CD82*** | 0.8000 | 0.0004 | 0.1030 | -0.0157 | 0.6410 | 0.0029 |
| ***TAP1*** | **0.0078** | 0.0413 | 0.1023 | 0.0158 | 0.1440 | 0.0282 |
| ***SMIM13*** | 0.0810 | 0.0180 | **0.0169** | 0.0335 | 0.1820 | 0.0237 |
| ***DDX3X*** | 0.0696 | 0.0009 | 0.9670 | < 0.001 | 0.6420 | 0.0029 |
| ***MYC*** | **0.0092** | 0.0397 | 0.710 | 0.0008 | 0.2290 | 0.0190 |
| ***APPL2*** | 0.3890 | 0.0044 | 0.610 | 0.0015 | 0.9700 | < 0.001 |

**Figure 5:**
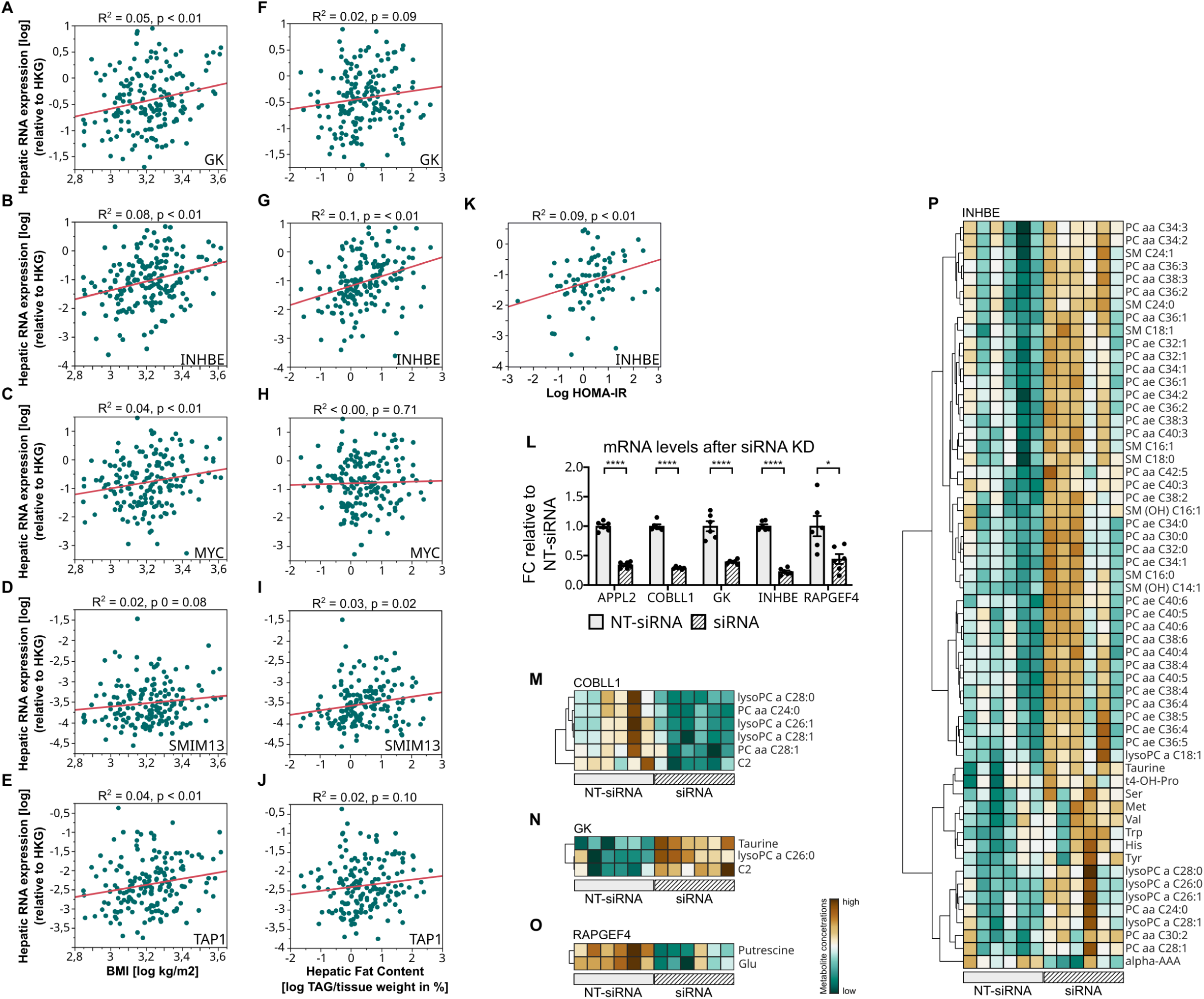
Networks genes are regulators of hepatic lipid metabolism. Pearson correlation analysis of human hepatic gene expression compared to BMI (log) **(A-E)** and hepatic TAG content in mg/100mg tissue (log) **(F-J)**, n = 170. (**K**) Human hepatic mRNA expression of INHBE correlated with HOMA-IR, n = 77. **(L)** Fold change (FC) in mRNA expression of five selected LRGs in HepG2 cells treated with siRNAs compared to non-targeting (NT-) siRNAs. Errorbars show standard error of the mean (SEM) (**M-P**) Metabolites significantly altered in HepG2 cells by the siRNA-mediated knockdown of four LRGs, compared to cells treated with NT-siRNA.

Finally, to test whether the expression of our candidate genes had an impact on cellular metabolite levels, we selected 5 representative genes to perform siRNA mediated knock-down (KD) experiments followed by metabolic profiling using the Absolute*IDQ*™ p180 Kit in HepG2 cells (Table S1). All five specific siRNAs significantly reduced target mRNA levels compared to a Non-Target-siRNA (Figure 5L). Subsequently, differences in metabolite concentrations for the specific siRNA KD samples were compared against their respective control sample using a one way ANOVA followed by Storey’s multiple testing correction [24]. For four of the five genes, namely *COBLL1, GK, RAPGEF4* and *INHBE*, we found between 2 and 58 metabolites to be significantly altered by siRNA mediated KD (Figure 5M-P). Overall, these data in HepG2 cells confirm a regulatory effect of the respective genes on the metabolic network, as predicted.

In summary, for seven of the eleven selected genes we could validate functional associations of human or murine hepatic transcript levels with parameters ranging from clinical obesity or insulin resistance in humans to changes in cellular metabolite levels. Of the three LRGs harboring SNPs that are associated to obesity or type 2 diabetes *COBLL1* and *INHBE* could be confirmed with at least one performed validation experiment, with human *INHBE* showing the strongest impact on cellular metabolism as it was significantly correlated with body weight, liver fat and whole-body insulin resistance.

## DISCUSSION

Inferring complex interactions of different types of biomolecules on multiple levels is a primary subject of systems biology. A steady increase in the generation of various omics data and the growing use of open science platforms and data repositories leads to a wealth of data that warrants further in-depth exploration. However, effective tools, especially for the integration of multi-omics data, are urgently needed to fully utilize this data gold mine. We here present the novel, fully unsupervised and data-driven method CoNI that allows for the integration of different omics data types based on a combined correlation approach followed by the construction of integrated hypergraphs. We successfully applied CoNI on two independent integration approaches. Primarily, we successfully integrated unpublished transcriptomics and metabolomics data from murine and human livers. Additionally, proteomics data combined with lipidomics from murine lungs were integrated to reconstruct known and potentially novel protein lipid interactions under clean air and smoking conditions (SI) [25, 26]. Thus, we postulate that our method is a versatile framework that can be applied to various data integration problems, not only including further omics data, but also including non-biological applications that aim to investigate a regulatory effect on network interactions.

Our new CoNI approach was able to uncover previously hidden and undescribed regulatory genes from liver metabolomics and transcriptomics data of chow- and HFD-fed mice that presumably play an important role in the development of liver steatosis in diet-induced obesity. By defining LRGs, we selected genes with a specific impact on a local sub-graph within the estimated networks. From the 79 LRGs identified, none was found to be present in both networks, illustrating that the genetic control of metabolic networks is widely altered by diet. Of note, only one of the genes selected for validation was among the 100 most significantly regulated genes between chow and HFD. Thus, the genes identified here would likely have been remained undiscovered by traditional approaches of analyzing RNA data alone. For five of the eleven further selected genes we found significant associations between human hepatic mRNA expression and major clinical features of metabolism, such as body weight, liver fat content and/or insulin resistance. Finally, with KD experiments in human HepG2 cells we could confirm four genes to significantly impact cellular metabolism. Taken together, seven of the eleven predicted genes could be experimentally validated. The protein-coding gene *Cobll1* had not been described in the context of HFD feeding, obesity and/or diabetes, but genetic variants of this gene are linked with type 2 diabetes related phenotypes. Here, we confirmed that *Cobll1* expression alters metabolite regulation in liver cells. Additionally, the SNP associated putative hepatokine *Inhbe was* positively confirmed in all performed validation experiments. This is in line with previous reports identifying *Inhbe* as diet-responsive gene in the rodent liver, regulated by HFD feeding, fasting or re-feeding [27-29]. Recently, Sugiyama et al. [23] demonstrated that the siRNA-mediated knockdown of *Inhbe* in obese insulin resistant Lep^db^ mice decreased fat mass and respiratory quotient thus suggesting enhanced whole-body fat utilization. We here link *Inhbe* to the regulation of AC, which are also known to interfere with hepatic insulin sensitivity [30, 31]. These two examples underline that the here presented method is a useful approach to successful integrate transcriptional data into metabolic networks to ultimately facilitate the identification of regulatory gene candidates of hepatic steatosis in mouse and humans. It can be used to integrate various types of multi-dimensional omics data and makes them available to useful holistic analyses in various fields of health research and beyond.

## METHODS

### Ethics Statement

*In vivo* experiments were performed without blinding of the investigators. All studies were based on power analyses to assure adequate sample sizes, and performed with approval by the State of Bavaria, Germany under the following protocol numbers: 55.2.1-54-2532-75-13. The clinical study has been approved by the Ethical review board of the University Hospital Tübingen and all human participants provided informed written consent.

### Animals

All experiments were performed in 20 adult male C57BL/6J mice purchased from Janvier Labs (Saint-Berthevin, Cedex, France). Mice were maintained on a 12-h light-dark cycle with free access to water and standard chow diet (Altromin, #1314). To promote diet-induced obesity (DIO), mice were ad-libitum fed with 58% high-fat diet (HFD) (Research Diets, D12331) for 22 weeks. Mice were fasted for 5 hours and then sacrificed by cervical dislocation for organ withdrawal. Livers were removed immediately, flash frozen in liquid nitrogen and stored at −80°C until further analysis. Two animals were excluded from the study cohort due to their comparatively lower body weight gain on HFD.

### Plasma analysis

Blood was collected in tubes containing 50 *µ*L EDTA and then centrifuged at 2000 x g and 4 °C for 10 min. Plasma was collected and stored at −80°C until further testing. Plasma triglycerides (TAG), cholesterol and non-esterified fatty acids (NEFA) were measured by commercial enzymatic assay kits (WAKO chemicals, Neuss, Germany). Insulin was measured by the ultrasensitive murine insulin ELISA kit (Merck Millipore, Darmstadt, Germany).

### Hepatic triglyceride content measurements

Hepatic triglyceride content was determined after chloroform/methanol (2:1) extraction by using the triglyceride assay kit according to the manufacturer’s protocol (Wako Chemicals).

### Metabolomics

#### Tissue Homogenization and Metabolite Extraction

Frozen murine liver samples were weighed and metabolites were extracted as previously described in ice-cold extraction solvent, a 85/15 (v/v) ethanol/10 mM phosphate buffer pH 7.5 mixture at a ratio of 3 *µ*L solvent per 1 mg tissue weight [32]. The liver samples were homogenized using a Precellys24 homogenizer (PeqLab Biotechnology, Erlangen, Germany) thrice for 20s at 5,500 rpm and 4°C, with 30 s pause intervals to ensure constant temperature, followed by centrifugation at 4°C and 10,000 x g for 5 min. Subsequently, 10 *µ*L of the supernatants were used for metabolite quantification.

#### siRNA Knockdown and Metabolite Extraction in HepG2 cells

Cells were cultured in DMEM supplemented with 10% fetal bovine serum and antibiotics (penicillin 100 IU/ml and streptomycin 100 *µ*g/ml) in 5% CO_2_ at 37 °C. At 70–80% confluence, cells were transfected with five different human SMARTpool On Target plus siRNA clones (L-006727-00, L-021435-02, L-009511-00, L-020477-02, L-016272-01; Dharmacon, Lafayette, USA) or ON-TARGETplus Non-targeting Pool (D-001810-10) using DharmaFECT #4 (Dharmacon) for 48 hours and subjected to metabolite extraction in an ice-cold 80/20 (v/v) methanol/water mixture as described previously [33]. Cells were homogenized using the Precellys24 homogenizer twice for 25 s at 5,500 rpm and 0-3°C, with 5s pause intervals in between. Subsequently, the samples were centrifuged at 4°C and 10,000 x g for 5 min and 20 *µ*L of the supernatants were used for the metabolite quantification. Each target specific siRNA and the Non-target control was transfected and measured in six biological replicates.

#### RNA extraction and qPCR in HepG2 cells

RNA was extracted from HepG2 cells after siRNA KD using the NucleoSpin RNA isolation kit (Macherey-Nagel, Düren, Germany). Equal amounts of RNA were reverse transcribed to cDNA using the QuantiTect Reverse Transcription kit (Qiagen, Hilden, Germany). Gene expression was analyzed using TaqMan probes for *APPL2* (Hs01565861_m1), *COBLL1* (Hs01117513_m1), *GK* (Hs02340007_g1), *INHBE* (Hs00368884_g1), *RAPGEF4* (Hs00199754_m1), and *HPRT* (Hs02800695_m1) as the housekeeping gene with the respective TaqMan mastermix (Thermo Fischer Scientific, Inc., Rockford, IL USA). qPCRs were carried out using a Quantstudio 6 Real Time PCR System (Applied Biosystems). Gene expression was evaluated using the Δ-Δ Ct method.

#### Fluorescence-based DNA quantification in cell homogenates

To normalize the obtained metabolomics data from cell homogenates for differences in cell numbers, the DNA content was determined using fluorochrome Hoechst 33342 (ThermoFisher Scientific, Schwerte, Germany) and a GloMax Multi Detection System (Promega, Mannheim, Germany) from a small aliquot taken before the final centrifugation step, as previously described [33]. The DNA content for one replicate of the *Cobll1* KD was too low, thus this sample was excluded from all downstream analysis, resulting in n=5 *Cobll1* KD replicates. **Metabolite Quantification by Absolute*IDQ*™ p180 Kit.** The targeted metabolomics approach was based on liquid chromatography-electrospray ionization-tandem mass spectrometry (LC-ESI-MS/MS) and flow injection-electrospray ionization-tandem mass spectrometry (FIA-ESI-MS/MS) measurements using the Absolute*IDQ*™ p180 Kit (BIOCRATES Life Sciences AG, Innsbruck, Austria). The assay allows simultaneous quantification of 188 metabolites out of 10 *µ*L tissue lysate or 20 *µ*L cell lysate, and includes free carnitine, 39 acylcarnitines (Cx:y), 21 amino acids (19 proteinogenic + citrulline + ornithine), 21 biogenic amines, hexoses (sum of hexoses – about 90-95 % glucose), 90 glycerophospholipids (14 lysophosphatidylcholines (LPC) and 76 phosphatidylcholines (PC)), and 15 sphingolipids (SMx:y). The abbreviations Cx:y are used to describe the total number of carbons and double bonds of all chains, respectively [32, 34]. For the LC-part, compound identification and quantification were based on scheduled multiple reaction monitoring measurements (sMRM). The method of Absolute*IDQ*™ p180 Kit has been proven to be in conformance with the EMEA-Guideline “Guideline on bioanalytical method validation (July 21st 2011)” [35], which implies proof of reproducibility within a given error range. Sample preparation and LC-MS/MS measurements were performed as described in the manufacturer manual UM-P180. The limits of detection (LOD) were set to three times the values of the zero samples (PBS). The lower and upper limit of quantification (LLOQ and ULOQ) were determined experimentally by Biocrates. The assay procedures of the Absolute*IDQ*™ p180 Kit as well as the metabolite nomenclature have been described in detail previously [36]. Metabolite concentrations were calculated using internal standards and reported in *µ*M.

### Transcriptomics

#### RNA preparation and Microarray analysis

Microarray data were obtained from liver samples of 10 chow and 9 HFD-fed mice. Total RNA was isolated from tissues employing a commercially available kit (NucleoSpin RNA, #740955, Macherey-Nagel, Düren, Germany). Total RNA (150 ng, RIN>7) was amplified using the WT PLUS Reagent Kit (Affymetrix, Santa Clara, US). Amplified cDNA was hybridized on Mouse Clariom S arrays (Affymetrix). Staining and scanning was done according to the Affymetrix expression protocol. Expression console (v.1.4.1.46, Affymetrix) was used for quality control and to obtain annotated normalized RMA gene-level data (Gene Level - SST-RMA). Genes with low expression levels (probe intensity < 40 in 5 out of 18 samples) were removed from the data set. For probe sets with identical values across all samples only one probe set was kept in the final gene sets. Before the calculation of the partial correlation coefficients, genes with high within-group variance (variance > 0.5) were excluded from the downstream analysis to reduce the number of identified false positives due to noisy expression patterns. This resulted in 10,159 gene expression profiles that were used in the downstream analysis.

### Human Data

#### Patients with liver tissue samples

For the analysis of gene expression in human liver tissue samples, a cohort of 170 men and women of European descendent undergoing liver surgery at the Department of General, Visceral, and Transplant Surgery at the University Hospital of Tübingen (Tübingen, Germany) was included in the present study. Subjects were fasted overnight before collection of the liver biopsies and in a subgroup of 77 individuals also fasting plasma samples for the calculation of the homeostasis model assessment of insulin resistance (HOMA-IR) were obtained as proposed by Matthews et al. [37]. Characteristics are shown in Table S5A for the whole group and in Table S5B for the subgroup with fasting plasma samples. All patients tested negative for viral hepatitis and had no liver cirrhosis. Only samples from normal, non-diseased tissue, judged by an experienced pathologist, were used. Informed, written consent was obtained from all participants, and the Ethics Committee of the University of Tübingen approved the protocol (239/2013BO1) according to the Declaration of Helsinki. Liver samples were taken from normal, non-diseased tissue during surgery, immediately frozen in liquid nitrogen, and stored at −80°C.

#### Determination of liver tissue triglyceride content

Liver tissue samples were homogenized in phosphate-buffered saline containing 1% Triton X-100 with a TissueLyser (Qiagen, Hilden, Germany). To determine the liver fat content, triglyceride concentrations in the homogenate were quantified using an ADVIA XPT clinical chemistry analyzer (Siemens Healthineers, Eschborn, Germany), and the results were calculated as TAG(mg)/100 mg tissue weight.

#### Real-Time PCR

For real-time (RT)-PCR and quantitative RT-PCR analyses of hepatic mRNA expression in liver biopsies, frozen tissue was homogenized in a TissueLyser (Qiagen), and RNA was extracted with the RNeasy Tissue Kit (Qiagen) according to the manufacturer’s instructions. Total RNA treated with RNase-free DNase I was transcribed into cDNA by using a first-strand cDNA kit, and PCRs were performed in duplicates on a LightCycler480 (Roche Diagnostics, Mannheim, Germany). The human primer sequences that were used are shown in Table S6. Data are presented relative to the housekeeping gene Rps13 using the Δ-Δ Ct method.

#### Quantification of blood parameters

Plasma insulin was determined on the ADVIA Centaur XPT chemiluminometric immunoassay system. Fasting plasma glucose concentrations were measured using the ADVIA XPT Clinical chemistry analyzer (both Siemens Healthineers, Eschborn, Germany).

### Data availability

The microarray data has been submitted to the GEO database at NCBI (GSE137923: https://www.ncbi.nlm.nih.gov/geo/query/acc.cgi?acc). All other data generated or analyzed during this study are available within the paper and its supplementary information file. Human data is not made available at the patient level due to data protection regulation.

### Statistics

#### Metabolite data analysis

After removal of metabolites with non-imputable concentration levels (> 5% missing values), TwoGroup from the R metabolomics package [38] was used to compare the log2-normalized concentration levels between the chow and HFD-fed mice. The siRNA KD altered metabolites in HepG2 cells were compared using a one way ANOVA with an FDR correction as introduced by Storey [24]. Unless otherwise stated, all metabolites showing a Benjamini-Hochberg [39] corrected p-value less than 0.05 were defined to differ significantly with respect to their concentration. For the downstream analysis the log2-transformed metabolite concentrations were scaled using the square root of the standard deviation as scaling factor (Pareto scaling) [40].

#### Differential gene expression analysis of murine liver samples

After log2-transformation, the R package limma [41] (Linear model for microarray data) was applied to infer differential expression between the two diet groups. We defined all genes with Benjamini-Hochberg corrected p-value less than 0.05 to be significantly deregulated.

#### Human data analyses

Data that were not normally distributed (Shapiro-Wilk W-test) were logarithmically transformed. Univariate associations between parameters were tested using Pearson correlation analyses. To adjust the effects of covariates and identify independent relationships, multivariate linear regression analyses were used. The statistical software package JMP 14.0 (SAS Institute, Cary, NC) was used.

#### Correlation analyses

For each metabolite pair *M*_*i*_ and *M*_*j*_ given *M, i = 1,…,n, j = 1,…,n, i*≠*j*, with *n* metabolites the Pearson correlation coefficients were obtained with R package Hmisc [https://CRAN.R-project.org/package=Hmisc]. Significant differences of metabolite correlations under different dietary conditions were tested using *Steiger’s test* [14] function of R’s cocor package [42].

#### Identification of main influencing factor

Principal component analysis (PCA) was performed on each set to find the main factor separating the samples. Tested factors were: Body weight measured at the end of the study (BWE), liver triglyceride level (TAG), and administered diet. To assess if the factors differ between chow and HFD we applied the Wilcoxon signed-rank test.

#### Pathway enrichment analysis of differentially expressed genes

Transcriptional enrichments were calculated using the R package ClusterProfiler [43] to test for overrepresented GO [44] biological process terms. We summarized terms that were completely contained in another term with respect to the enriched gene list.

#### Partial Correlation based Network Integration (CoNI)

The framework (Figure 1) includes three steps carried out for each treatment group independently: 1) performing pairwise correlation analysis on metabolite data set; 2) Partial correlation analysis combining the metabolite concentrations with the gene expression profiles; and 3) Construction of undirected, weighted graph.

##### 1) Correlation analysis

First, for *M* metabolites the *M*x*M* correlation matrix was calculated. Here, for subsequent analyses only metabolite pairs showing a Pearson correlation p-value < 0.05 were selected.

##### 2) Partial correlation based gene extension

For each pair of selected metabolites M_i_ and M_j_ given *M, i = 1,…,M, j = 1,…,M, i*≠*j* and each gene G*k* with *k = 1,…,K* the partial correlation *ρ(M*_*i*_*M*_*j*_*\*G*_*k*_) reflecting the correlation between *M*_*i*_ and *M*_*j*_ after removing the linear effects of gene *G*_*k*_ were calculated using R’s package ppcor [45]. Such a combination was denoted as triplet in the following. Steiger’s test [14] was adapted to select triplets, where the partial correlation coefficient differed significantly from the correlation coefficient of the respective metabolite pair. The original test assesses the significance for the difference between two correlation coefficients that have one variable in common. The significance depends on the intercorrelation between the two variables that are not shared, which has to be provided as additional parameter. To be able to use this test and compare a partial correlation coefficient and a correlation coefficient, the test was applied twice. The provided, additional parameter was in the first test the correlation between M_*i*_ and G*k* and in the second test the correlation between M*j* and G*k*. To be selected, the triplet had to significantly reject the null-hypothesis (Bonferroni adjusted p-value < 0.05), stating that the correlation coefficients did not differ in both tests. The method cocor.dep.groups.overlap of R’s cocor package [42] was used to perform the testing.

##### 3) Undirected graph construction and clustering

Next, an un-directed and weighted graph was generated where nodes are formed by metabolites and genes set up the edges. Edges were drawn if a metabolite pair correlated and this correlation was significantly influenced by at least one gene. Several genes can connect more than two correlating metabolites and a pair of metabolites might also be connected by more than one gene. The number of genes connecting the two respective metabolite nodes determines the edge weight.

#### Local regulator genes (LRGs)

Starting from each node in the network, the appearance of each gene in all edges connecting nodes with a distance <= two was counted. Statistical significance was estimated using a binomial distribution test. P-values were Bonferroni corrected for multiple testing and genes with adjusted p-value < 0.05 were defined as LRGs.

#### Communities

To find densely connected sub-graphs in the graph, the fast greedy modularity optimization algorithm [46] implemented in the igraph package [47] was applied.

## Funding

This work was supported in part by a grant from the German Federal Ministry of Education and Research (BMBF) to the German Center for Diabetes Research (DZD e.V.), by the Helmholtz Portfolio Program “Metabolic Dysfunction” (MHT), by the Alexander von Humboldt Foundation (MHT), by DZD tandem grant funds (SCS, PP, MH), by the Helmholtz-Israel-Cooperation in Personalized Medicine (PP), by the Helmholtz Initiative for Personalized Medicine (iMed; MHT), by Helmholtz Alliance Aging and Metabolic Programming (AMPro) and through the Initiative and Networking Fund of the Helmholtz Association.

## Acknowledgments

We gratefully acknowledge Alke Guirguis and Ann-Kathrin Horlacher (Institute for Clinical Chemistry and Pathobiochemistry, University Hospital Tübingen, Germany) and Silke Becker (Research Unit Molecular Endocrinology and Metabolism, HMGU) for their technical assistance.

## Author Contributions

V.S.K. and D.L. conceived the method. V.S.K. and J.M.M-K. wrote the code and analyzed the data. S.C.S. performed the mouse study and experiments. M.I. and J.B. performed transcriptomics microarray analysis. J.T., C.P., G.K. and J.A. performed metabolomics. A.P., A.K. and M.H. contributed the human liver data. M.H.T. and T.D.M. contributed reagents, materials and critical input. P.T.P. and D.L. conceptualized the project. V.S.K., S.C.S., P.T.P. and D.L. interpreted the data. V.S.K., S.C.S. and D.L. and wrote the manuscript. All authors critically reviewed and edited the manuscript and approved the final version.

## Competing Interests

Matthias H. Tschöp is a scientific advisor to Novo Nordisk, and ERX. Jerzy Adamski is a scientific advisor to Biocrates Life Sciences AG.

## Notes

### Competing Interest Statement

Matthias H. Tschoep is a scientific advisor to Novo Nordisk, and ERX. Jerzy Adamski is a scientific advisor to Biocrates Life Sciences AG.

