## Supplementary material for "Correlation guided Network Integration (CoNI) reveals novel genetic regulators of hepatic metabolism": Suplementary Figures Tables and Information

**SUPPLEMENTARY INFORMATION**

### Figure S1

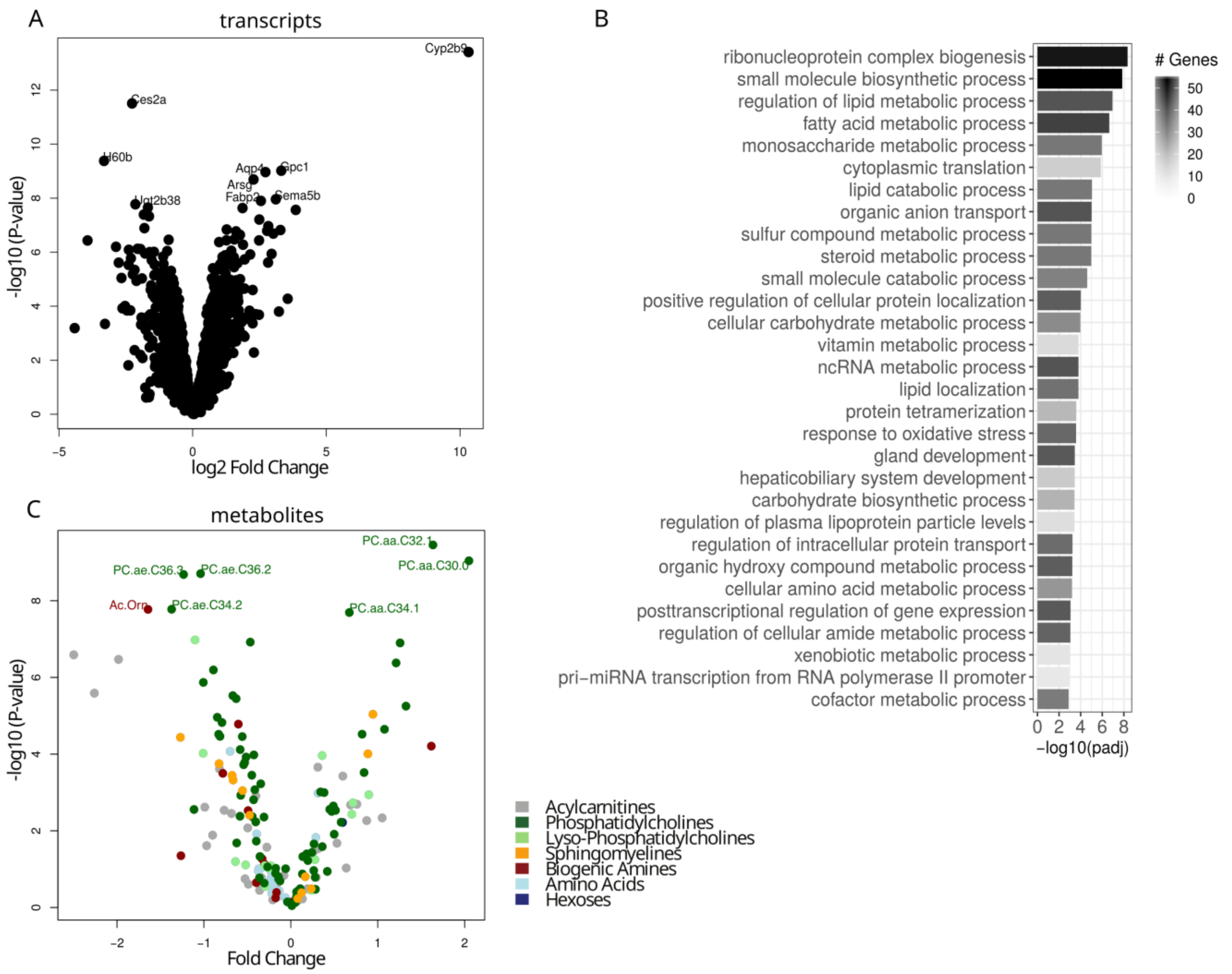


**Supplementary Figure S1: Differential analysis of RNA and metabolites. (A)** Volcano plot of hepatic transcript expression. **(B)** Gene Ontology (Biological Process) enrichment of significantly regulated genes (p<0.05, Benjamini Hochberg corrected). Redundant terms were removed from this list (Full List available as Table S2). **(C)** Volcano plot of hepatic metabolite regulation. Metabolite classes are indicated by colors.

**Figure S2**

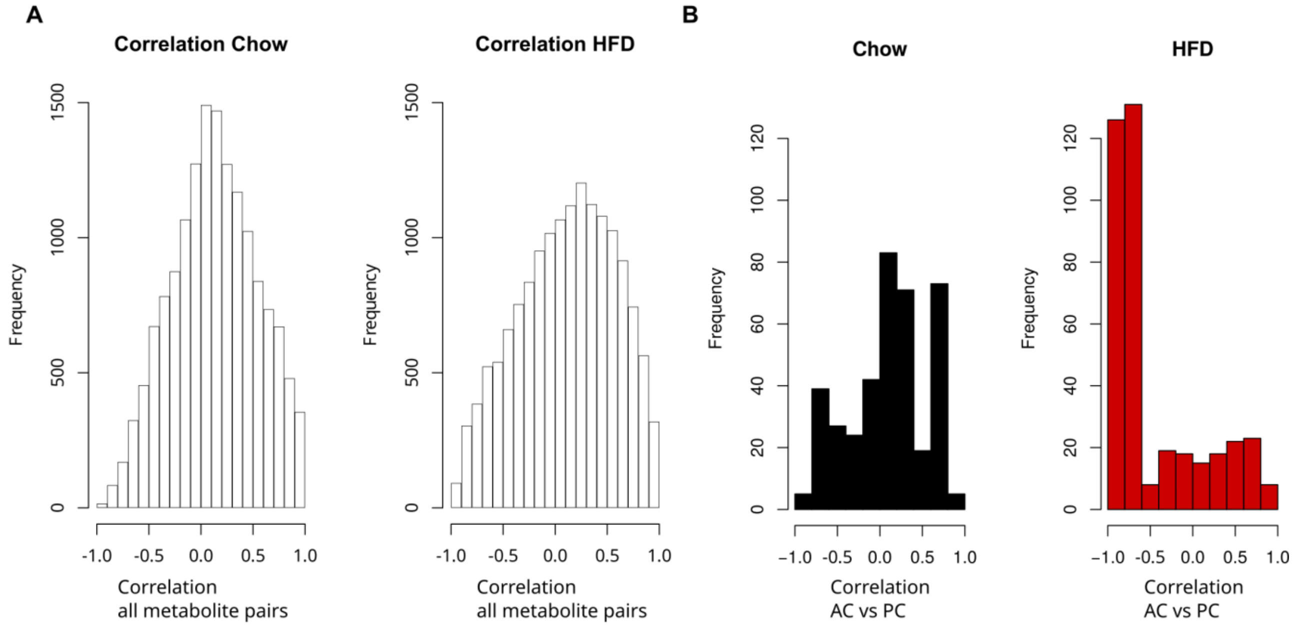


**Supplementary Figure S2: Correlation comparison. (A)** Distribution of Pearson's correlation coefficients between all pairs of metabolites under chow and HFD. **(B)** Distribution of Pearson's correlation coefficients between pairs of acylcarnitines and phosphatidylcholines under chow and HFD.

**Figure S3**

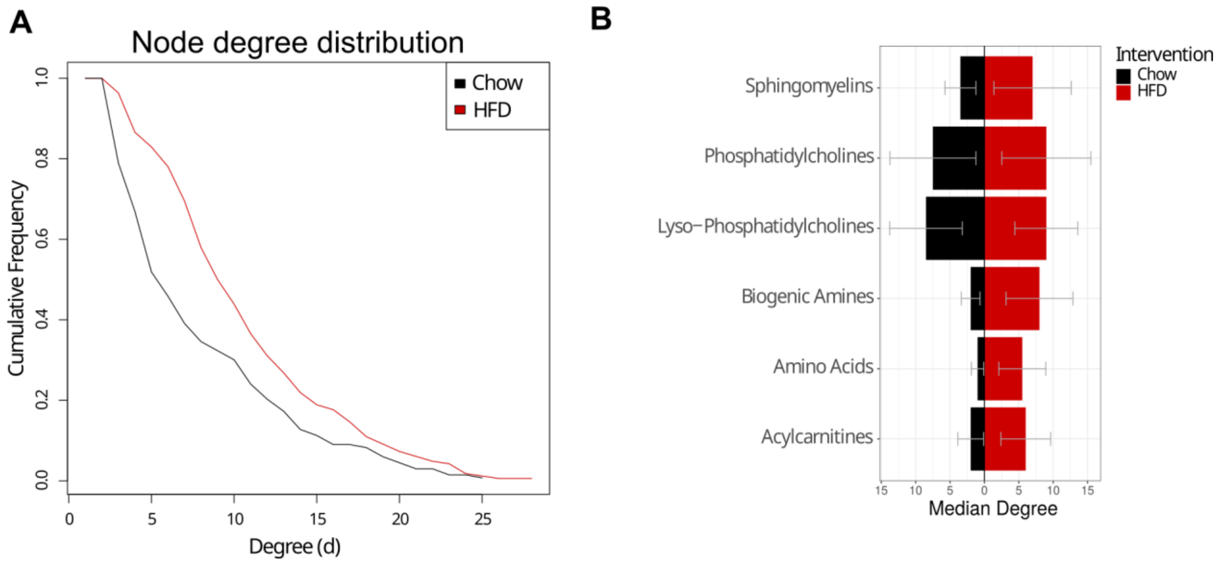


**Supplementary Figure S3: Node degree comparison. (A)** Node degree distribution in the integrated networks for chow (black) and HFD (red). **(B)** Node degree distributions for metabolite classes compared between chow and HFD. Errorbars show the standard deviation for each metabolite class.

**Figure S4**

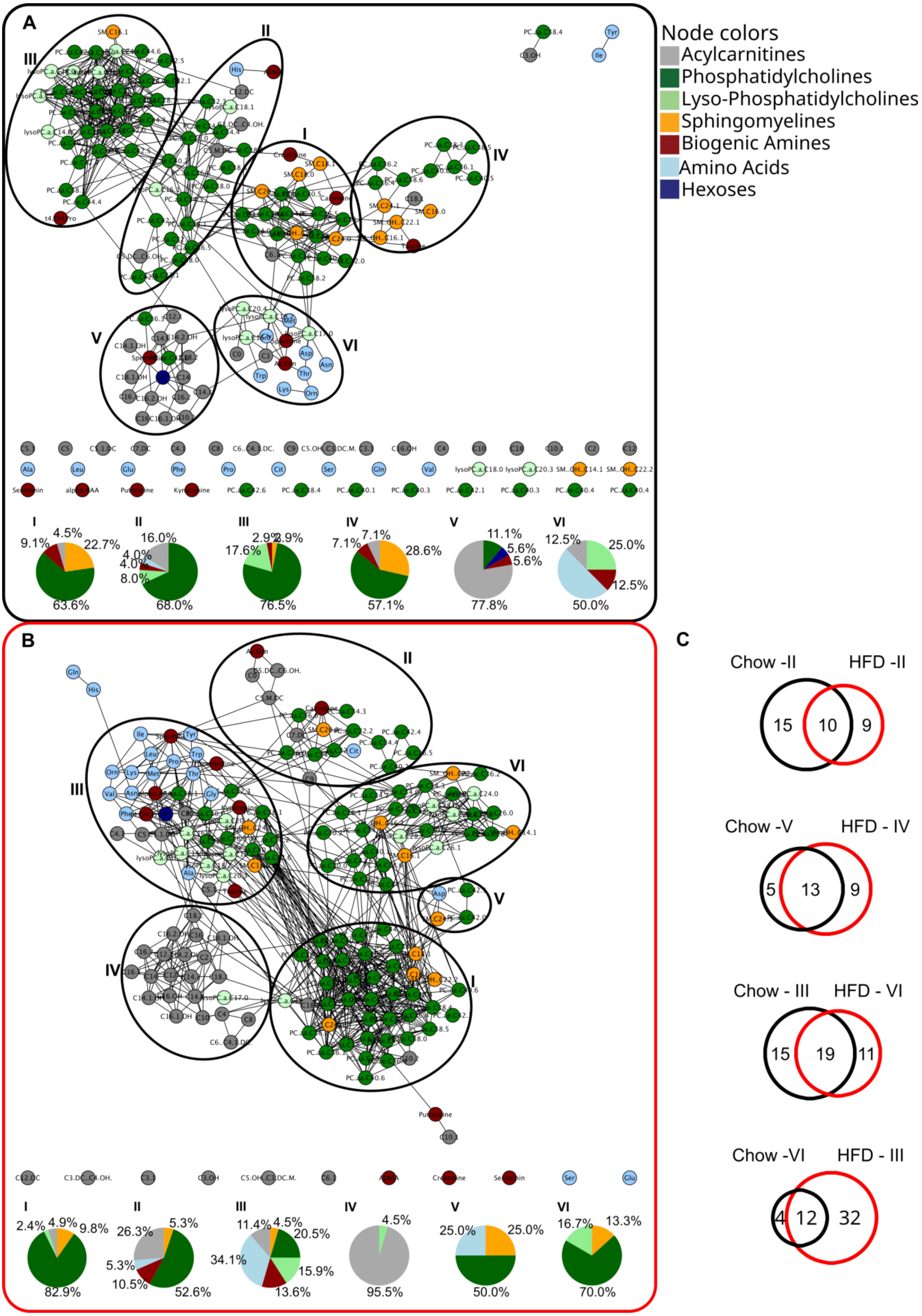
A
Node colors
Acylcarnitines
Phosphatidylcholines
Lyso-Phosphatidylcholines
Sphingomyelines
Biogenic Amines
Amino Acids
Hexoses
I
4.5%
9.1%
22.7%
63.6%
II
16.0%
4.0%
4.0%
8.0%
68.0%
III
2.9%
2.9%
17.6%
76.5%
IV
7.1%
7.1%
28.6%
57.1%
V
11.1%
5.6%
5.6%
77.8%
VI
12.5%
25.0%
12.5%
50.0%
B
I
2.4%
4.9%
9.8%
82.9%
II
26.3%
5.3%
5.3%
10.5%
52.6%
III
11.4%
4.5%
20.5%
34.1%
15.9%
13.6%
IV
4.5%
95.5%
V
25.0%
25.0%
50.0%
VI
16.7%
13.3%
70.0%
C
Chow -II
HFD -II
15
10
9
Chow -V
HFD - IV
5
13
9
Chow - III
HFD - VI
15
19
11
Chow -VI
HFD - III
4
12
32

**Supplementary Figure S4: Network Metabolites are organized in communities.** Inferred networks for chow **(A)** and HFD **(B)**. Both networks tend to be organized in communities. Densely connected sub-networks indicated by circles are labeled in Latin numbers (I - VI). They were identified applying the fast greedy modularity optimization algorithm [45]. Metabolite composition of each community is summarized in the pie charts below the networks. **(C)** Overlaps of metabolite communities between chow and HFD are compared. For both diets the PC split up in four communities whereas the other two divide up in one with mainly AC and one mainly build by AA. Whereas the latter communities (AC and AA) appear to be stable, the PC seem to be more reorganized upon diet change.

**Figure S5**

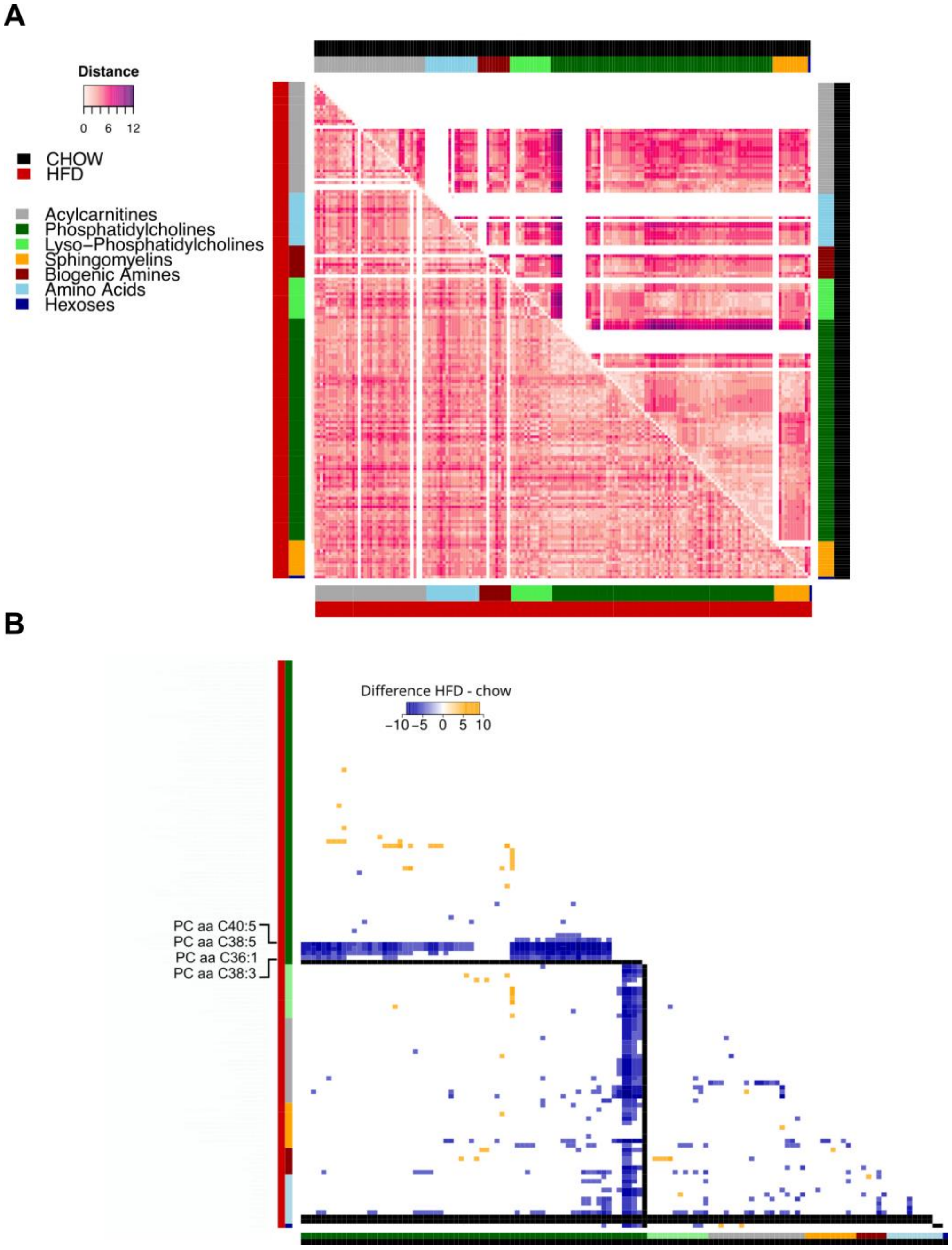


**Supplementary Figure S5: Network shortest path distances. (A)** The reorganization of metabolites in the two graphs from chow- and HFD-fed mice was compared by the minimal distances (path length) between all pairs of metabolites. Non-connected metabolites are uncolored. In general, distances in chow were higher than in the HFD network. **(B)** Changes in path length from chow to HFD are displayed as differences**.** The most striking differences were observed for the four PC PCaaC38:3, PCaaC36:1, PCaaC40:5, and PCaaC38:5.

**Figure S6**

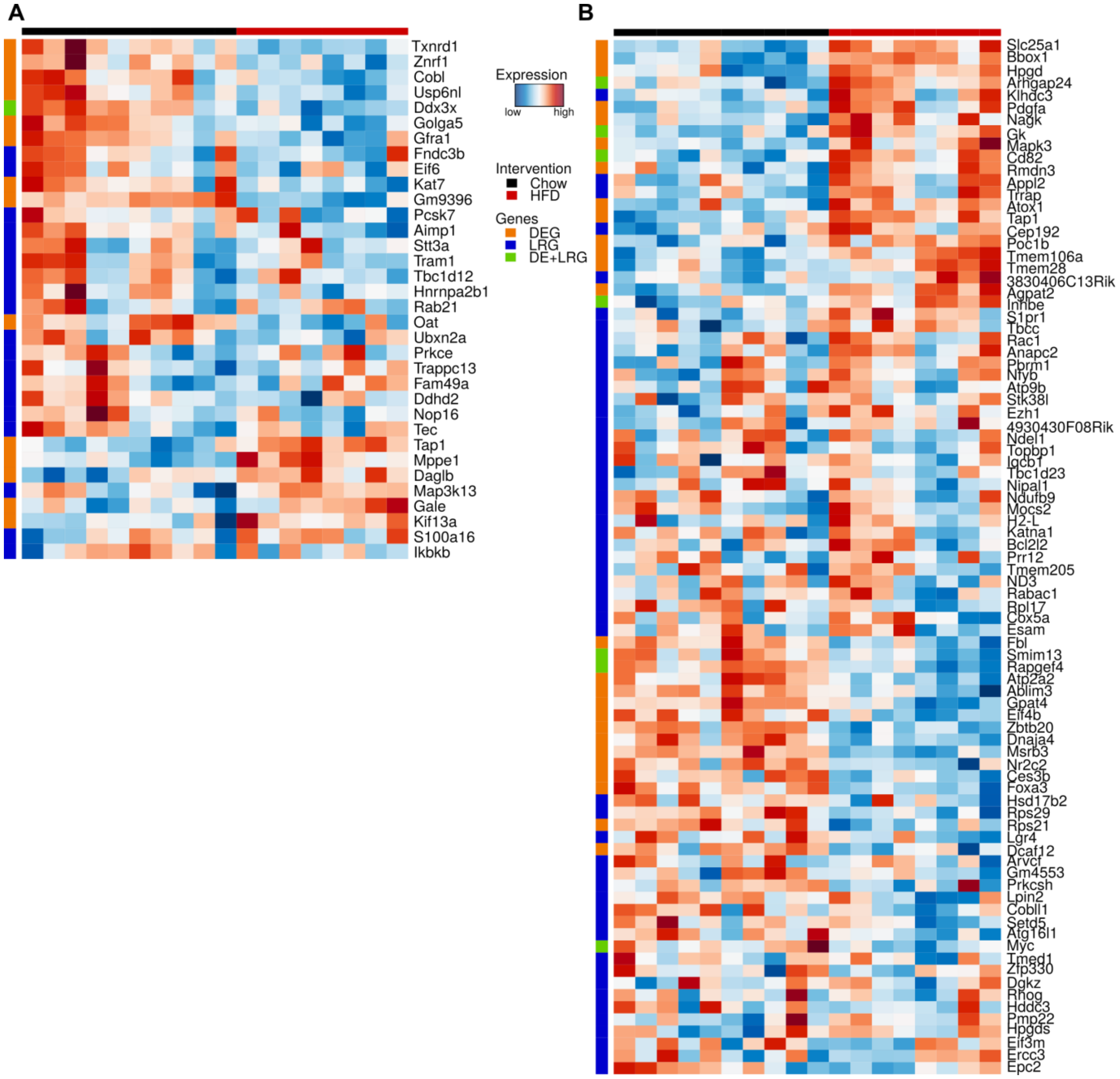


**Supplementary Figure S6: Gene expression pattern of network genes.** For chow (**A**) and HFD (**B**) network genes are shown that are either identified as LRG (blue, left color bar), differentially expressed (orange, left color bar), or both (green, left color bar).

### Figure S7

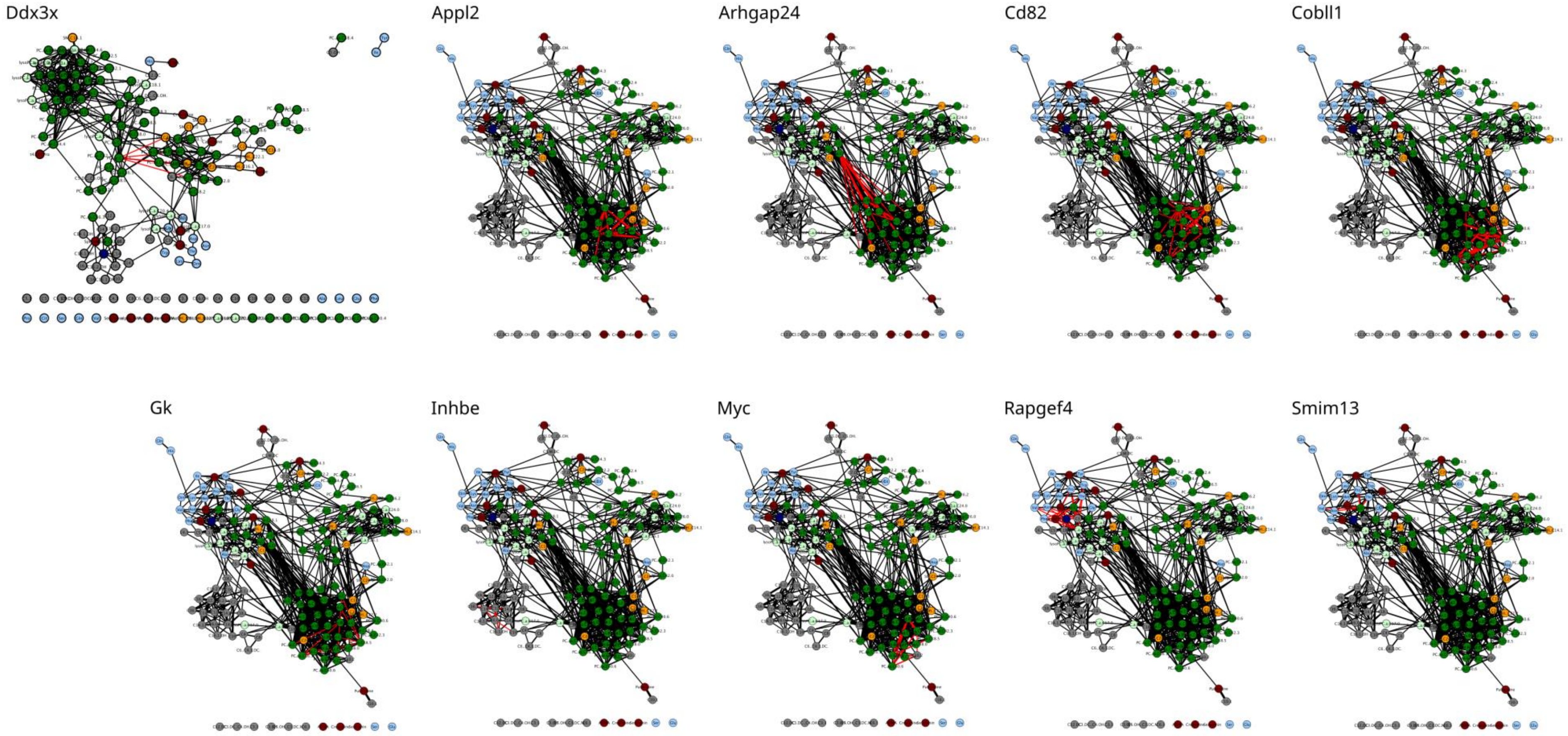


**Supplementary Figure S7: Network distributions of selected LRGs.** Edges containing the respective LRG are colored red

### Figure S8

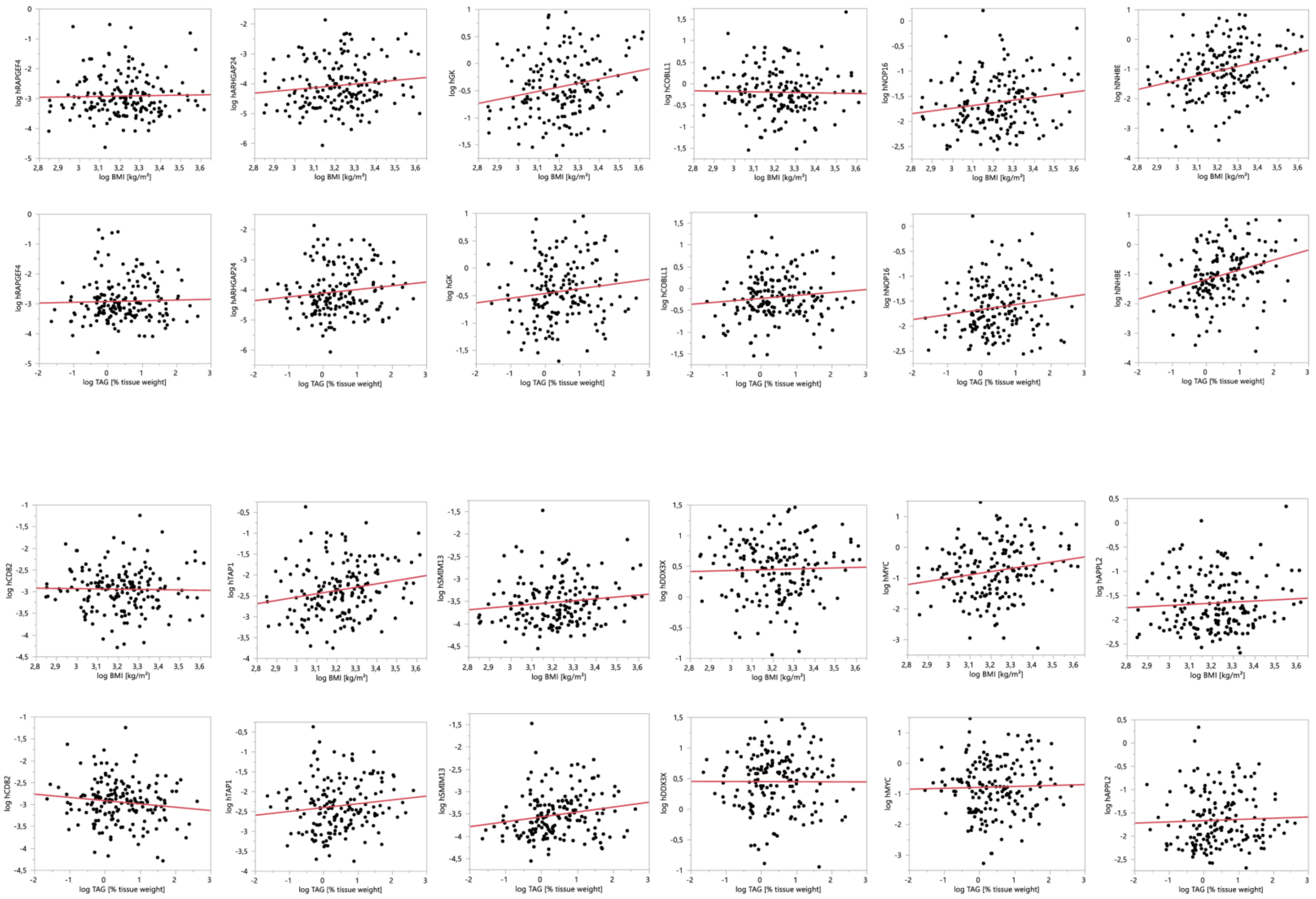


**Supplementary Figure S8:** Pearson correlation analysis of human hepatic gene expression compared to BMI (log) and hepatic TAG content in mg/100mg tissue (log)

### Supplementary Table S4

Identified SNPs associated with obesity, insulin resistance and type 2 diabetes-related phenotypes. SNPs were identified using the Type 2 Diabetes Knowledge Portal. Only phenotypes with a strong ($p < 0.05$) association for variants within the genes were selected.

| | **BMI** | **Waist-hip ratio** | **Waist-hip ratio adj BMI** | **Waist circumference** | **Hip circumference** | **Triglycerides** | **Cholesterol** | **HDL cholesterol** | **LDL cholesterol** | **Fasting insulin** | **Type 2 diabetes** | **Type 2 diabetes adj BMI** | **eGFR-creat (serum creatinine)** | **Heart rate** | **Incremental insulin at 30 min OGTT** | **HOMA-B** | **HbA1c** | **Creatinine** | **Systolic blood pressure** | **Diastolic blood pressure** | **Coronary artery disease** |
|---|---|---|---|---|---|---|---|---|---|---|---|---|---|---|---|---|---|---|---|---|---|
| ***Cobll1*** | x | x | x | x | x | x | | x | x | x | x | x | | | | | | | | | |
| ***Appl2*** | x | x | | x | | | | | | | x | | x | x | x | x | x | | | | |
| ***Topbp1*** | x | | | x | | | | | | | x | | | | | | | x | x | x | |
| ***Pbrm1*** | x | x | x | x | | | | x | | | | | | | | | | | | | |
| ***Lgr4*** | x | | | x | x | | | | | | | | | | | | | | | | |
| ***Tmed1*** | | | | | | | x | | x | | | | | | | | | | | | x |
| ***Rpl17*** | | | | | | | x | x | | | | | | | | | | | | | |
| ***Atp9b*** | x | | | | | | | | | | | | x | | | | | | | | |
| ***Rac1*** | x | | | | | | | x | | | | | | | | | | | | | |
| ***Tmem205*** | | | | | | | x | | x | | | | | | | | | | | | |
| ***Hddc3*** | | | | | | | | | | | x | x | | | | | | | | | |
| ***Inhbe*** | | | | | | x | | x | | | | | | | | | | | | | |
| ***Dgkz*** | | | | | | | | x | | | | | | | | | | | | | |
| ***Cep192*** | x | | | | | | | | | | | | | | | | | | | | |
| ***Tbcc*** | x | | | | | | | | | | | | | | | | | | | | |
| ***Setd5*** | x | | | | | | | | | | | | | | | | | | | | |
| ***Cox5a*** | x | | | | | | | | | | | | | | | | | | | | |

**Supplementary Table S5A**

Characteristics of subjects who donated liver samples

| | N | Means ± SD | Median (25–75%) |
|---|---|---|---|
| N, Gender (males/females) | 170 (106/64) | | |
| Age, years | | 63.5 ± 11.7 | 65 (58–72) |
| Body mass index, kg/m2 | | 25.2 ± 4.1 | 25.2 (22.2–27.5) |
| Liver fat content, % | | 2.16 ± 2.08 | 1.36 (0.85–2.65) |

**Supplementary Table S5B**

Characteristics of subjects with available fasting blood samples

| | N | Means ± SD | Median (25–75%) |
|---|---|---|---|
| N, Gender (males/females) | 77 (49/28) | | |
| Age, years | | 62.1 ± 12.1 | 64 (55–71) |
| Body mass index, kg/m2 | | 24.7 ± 4.1 | 24.3 (21.5–26.7) |
| Liver fat content, % | | 1.72 ± 1.35 | 1.22 (0.85–2.11) |
| Fasting glucose, mg/dl | | 98.5 ± 25.8 | 92 (82–115) |
| HOMA-IR, AU | | 2.26 ± 2.96 | 1.38 (0.71–2.36) |

HOMA-IR, homeostasis model assessment of insulin resistance; AU, arbitrary units.

**Supplementary Table S6**

Human primer sequences for real-time PCR

| **Human Gene** | **Upstream Primer** | **Downstream Primer** |
|---|---|---|
| RPS13 | 5′-ccccacttggttgaagttga-3′ | 5′-acaccatgtgaatctctcagga-3 |
| ARHGAP24 | 5′-tgtcttgagctcccagcaa-3′ | 5′-tgacaaagcctccttgcttc-3′ |
| CD82 | 5′-gccgacaagagcagtttcat-3′ | 5′-gacataggcccccatccta-3′ |
| GK | 5′-ttgattcatggcttatttggagt-3′ | 5′-tctgtacagtggacacctccat-3′ |
| INHBE | 5′-tcagctttgctactgtcacagac-3′ | 5′-cgaggagtggacaggtgaa-3′ |
| MYC | 5′-tgctccatgaggagacacc-3′ | 5′-cttttccacagaaacaacatcg-3′ |
| RAPGEF4 | 5′-ttttatgccaaatacccagctt-3′ | 5′-tgaaggctgtgcgtggta-3′ |
| SMIM13 | 5′-ctgactctgcttgtgttcgtg-3′ | 5′-agatgccatacaaaataccaacc-3′ |
| DDX3X | 5′-gctggcctagacctgaactc-3′ | 5′-gcttctcggttccttaaatgag-3′ |
| COBLL1 | 5′-ccgagtcacctagtgccagt-3′ | 5′-ttcattatgtgcagagttattttcct-3′ |
| APPL2 | 5′-caagcagtgactcccattacaa-3′ | 5′-tcattttcattttccatctctgaa-3′ |
| TAP1 | 5′-ctcagggctatgacacagagg-3′ | 5′-acacggtttccggatcaat-3′ |

**Additional Discussion of Selected Network Genes**

For the following genes, we found further evidence to be involved in lipid metabolism or diabetic related topics. The membrane-associated protein *Tap1* that was the only non LRG but present in both networks was recently linked to the initiation and propagation of liver inflammation as well as insulin resistance in mice [46], which is in line our association of hepatic TAP1 expression linked to BMI in humans. The adaptor protein *Appl2* has been shown to be involved in insulin signaling, endosomal trafficking, adiponectin signaling and other signaling pathways [47]. Ryu [48] demonstrated that a liver-specific knockout of *Appl2* improved insulin sensitivity, increased adiponectin signaling, and induced anti-inflammatory effects in HFD-fed mice, implicating the involvement of *Appl2* in hepatic metabolism. The LRG glycerol kinase (*Gk*) that had been proposed as regulator for several lipids was upregulated on HFD suggesting an adaptive mechanism to handle the increased hepatic lipid load. This hypothesis is supported by the finding that overexpression of *Gk* favours recycling of free fatty acids leading to increased fat storage in rat hepatoma cells [49-52]. Also *Myc* seems to be involved in the regulation of hepatic glycolysis [53-55]. Under HFD exposure, *Myc* overexpression in transgenic mice normalises glycemia, insulinemia, and the expression of genes involved in hepatic metabolism [54].

The six remaining LRGs identified (*Arhgap24, Cobll1, Cd82, Ddx3x, Smim13, Rapgef4*) have so far not been discussed in the context of obesity related hepatic liver metabolism.

**Application of CoNI for the integration of proteomics and lipidomics data**

To further test CoNI on a second publically available dataset, we used a combined lipidomics and proteomics dataset published by Phillips et al. and Titz et al. [56, 57]. In these studies, the effect of smoke from modified risk tobacco products on the lungs was tested in mice. For our purposes, we used the available protein and lipid data of mice exposed to cigarette smoke (CS), and mice exposed to fresh air (control). The normalized protein data was used as published and the lipid data was log2-transformed. Missing data from both datasets were filtered out (without imputation). When correlating only metabolites, we identified 571 significantly correlated lipid pairs for the control and 660 for CS (non-adjusted $p < 0.05$). Subsequently by applying CoNI we generated two networks (Figure S8A, B) consisting of 2,613 triplets (protein and lipid pairs) for the control and 1,973 triplets for CS. Of the 204 lipids used in the analysis, 116 and 112 lipids were present in the control and CS networks, respectively (Figure S8A, B). From these lipids, five were unique in the control and one in the CS-network. From the connected lipid pairs in the networks, 81 were shared between the two networks, 277 were unique for the control and 426 for CS. The CS-network showed a higher node degree distribution compared to the control (Figure S9). In the control network 117 local regulator proteins (LRPs) were identified and 82 for CS. Only three LRPs were shared between treatments (Pcmt1, Hsd11b1, Ltf).

To further review the CoNI delivered integrated proteins, we compared our findings to the results reported by Phillips et al. and Titz et al. [56, 57]. From the 82 LRP identified in the CS network, eleven were discussed in either one of the articles (Figure S10). Among those were core fatty acid synthase (Fasn), surfactant protein C (Sftpc) and carbonyl reductase 3 (Cbr3). Cbr3 is involved in the arachidonic acid metabolism pathway [57] and catalyzes the reduction of many endogenous and xenobiotic carbonylcompounds, including steroids and prostaglandins [58]. Sftpc was identified to regulate fatty acids and eicosanoids (Figure S10).

Sftpc is a major structural component of the lung surfactant, which is an important lipid-protein complex for lung homeostasis [57] and mutations in this gene have been linked with interstitial lung disease [59]. Under CS conditions, we identified Fasn to mainly regulate the sphingolipids Cerebroside/Ceramides (Figure S10) which are increased in CS-exposed mice and associated to lung cancer [60, 61]. Fasn is known to be upregulated in smokers [62] and is involved in fatty acid biosynthesis, which is important for glycerolipids and glycerophospholipids metabolism in the context of de novo synthesis of lipids due to CS-induced damage [56, 63]. Furthermore, we found a conspicuous clustering of ceramides and cerebrosides/glycosylceramides in the center of the CS network (Figures S8B, S10). CS induces the production of ceramides, which are involved in inflammation as well as alveolar destruction [57], lung endothelial cell apoptosis [64] and induction of metabolic disruption in mice [65]. With the CoNI approach, we here identified two LRPs that potentially regulate these ceramides, namely Fasn and the Membrane Metalloendopeptidase Mme. Whereas Fasn is known and frequently discussed to be induced by CS, Mme is not known to be involved in CS conditions, yet. These results additionally show that CoNI is a powerful method to integrate omics-data and capable of capturing relevant biological signals and also to identify so far unknown and possible target genes and proteins.

**Figure S8**

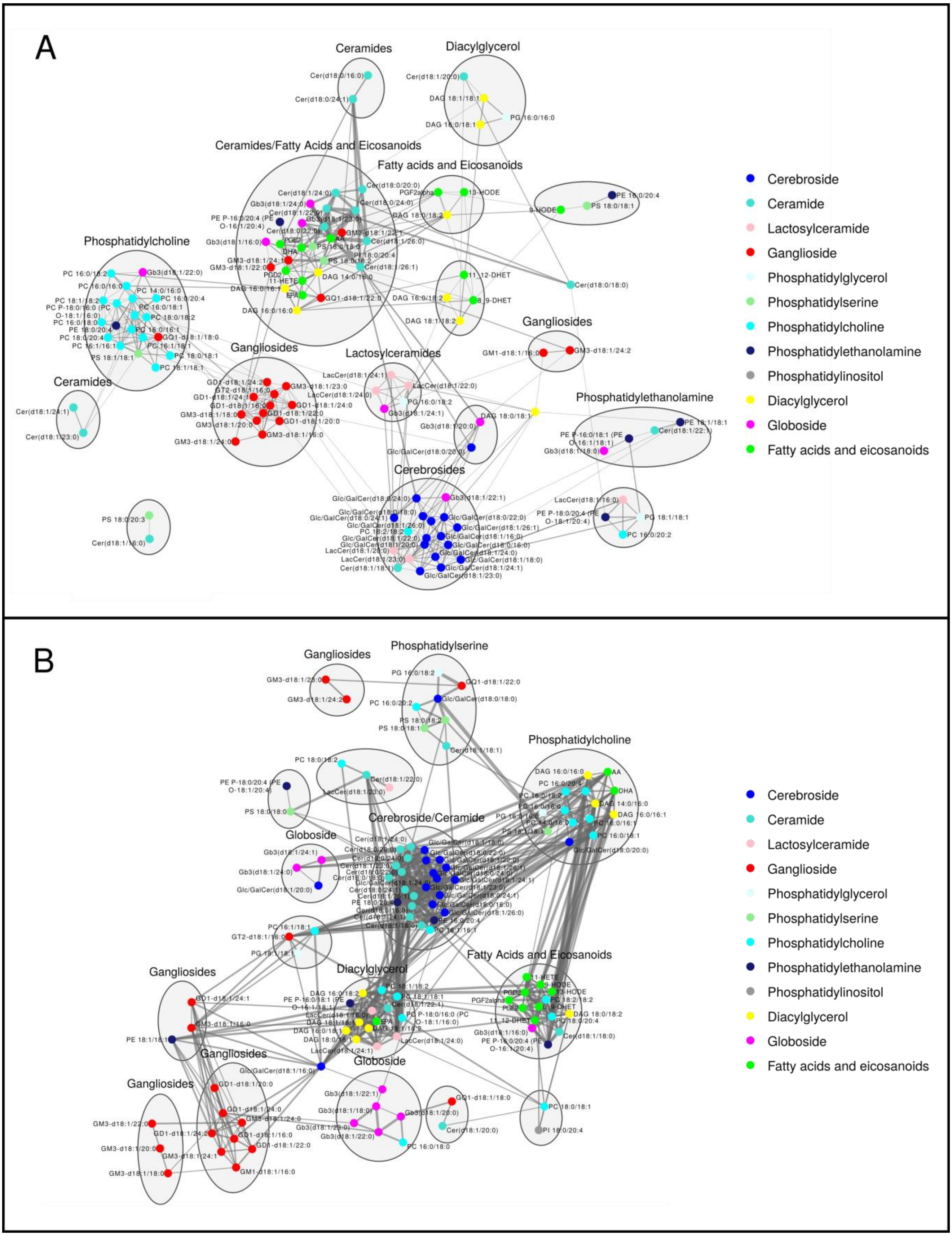
A
Ceramides
Diacylglycerol
Ceramides/Fatty Acids and Eicosanoids
Fatty acids and Eicosanoids
Phosphatidylcholine
Gangliosides
Lactosylceramides
Gangliosides
Ceramides
Phosphatidylethanolamine
Cerebrosides
Cerebroside
Ceramide
Lactosylceramide
Ganglioside
Phosphatidylglycerol
Phosphatidylserine
Phosphatidylcholine
Phosphatidylethanolamine
Phosphatidylinositol
Diacylglycerol
Globoside
Fatty acids and eicosanoids
B
Gangliosides
Phosphatidylserine
Phosphatidylcholine
Globoside
Cerebroside/Ceramide
Diacylglycerol
Fatty Acids and Eicosanoids
Gangliosides
Gangliosides
Globoside
Gangliosides
Cerebroside
Ceramide
Lactosylceramide
Ganglioside
Phosphatidylglycerol
Phosphatidylserine
Phosphatidylcholine
Phosphatidylethanolamine
Phosphatidylinositol
Diacylglycerol
Globoside
Fatty acids and eicosanoids

**Supplementary Figure S8: Integrated graphs of murine lung lipidome and proteome data generated using CoNI.** Networks generated using CoNI with lung lipid and protein data of mice exposed to fresh air (A) and cigarette smoke (B). The log2 of the concentrations of the lipid data was calculated and used for running CoNI. Cytoscape (v3.7.2) was used for network visualization and lipids were clustered using the MCL clustering algorithm tool, clusters are indicated with areas. Labels for the clusters correspond to the most abundant lipid group.  Thickness of the edges is proportional to the number of proteins affecting the respective lipid pair. Node color refer to lipid classes.

**Figure S9**

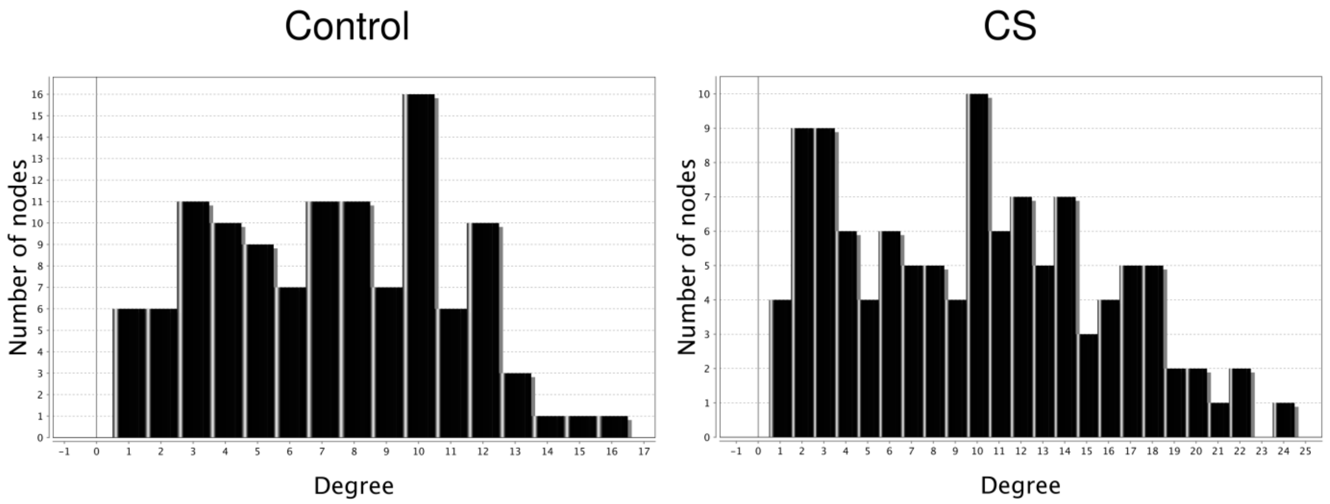


**Supplementary Figure S9: Degree distributions for the two estimated networks.**

**Figure S10**

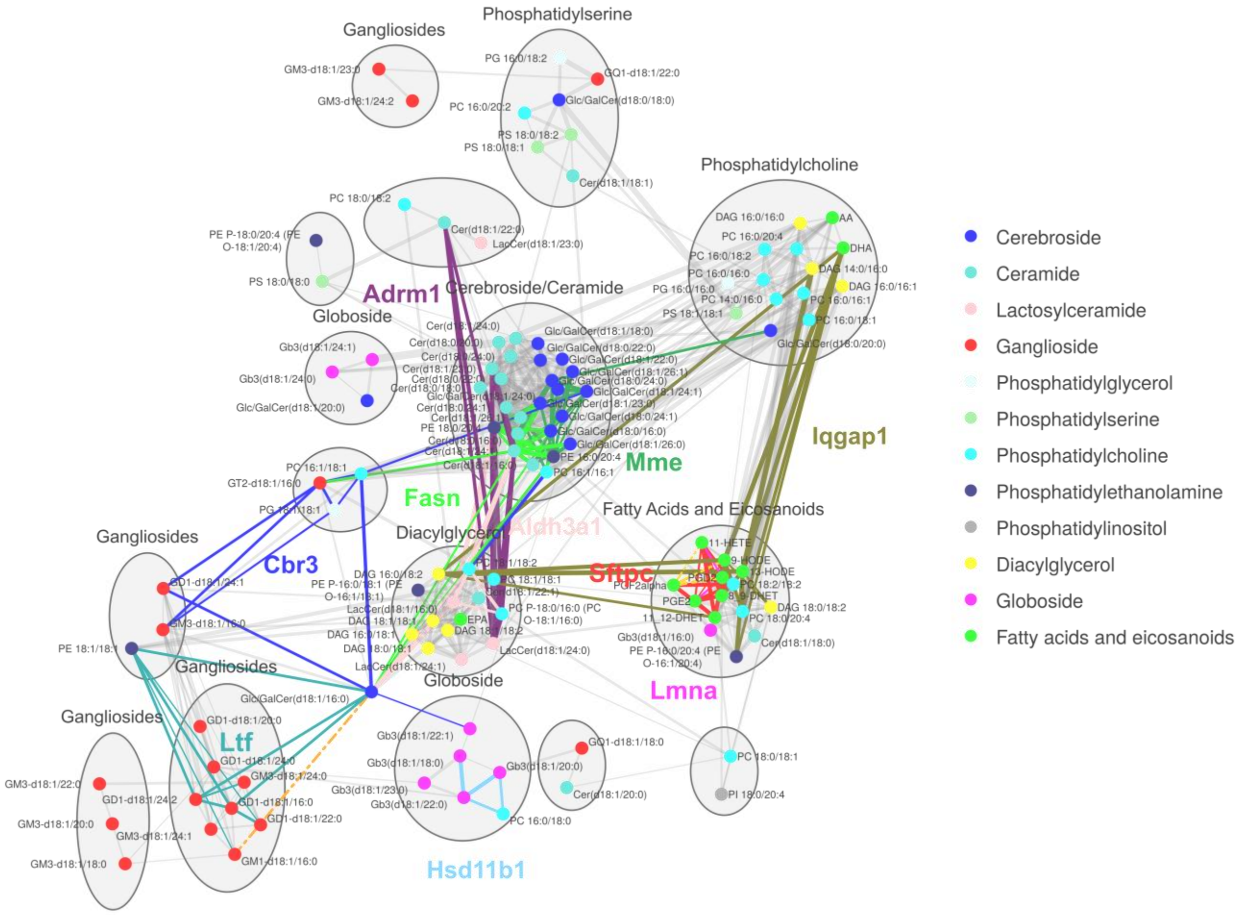


**Supplementary Figure S10: CoNI network of mice exposed to cigarette smoke.** Edges containing local regulator proteins relevant to the effects of smoking in the lungs and discussed in the publications of Philips et al. [56] and Titz et al. [57] are marked in different colors. Cytoscape (v3.7.2) was used for network visualization and lipids were clustered using the MCL clustering algorithm tool, clusters are indicated with areas. Labels for the clusters correspond to the most abundant lipid group. Thickness of the edges is proportional to the number of proteins affecting the respective lipid pair. Node Colors refer to lipid classes.
